# Fluorescence based live cell imaging identifies exon 14 skipped hepatocyte growth factor receptor (MET) degraders

**DOI:** 10.1101/2024.11.22.624922

**Authors:** Jayapal Reddy Mallareddy, Lin Yang, Wan-Hsin Lin, Ryan Feathers, Jennifer Ayers-Ringler, Ezequiel Tolosa, Amritha G. Kizhake, Smitha Kizhake, Sydney P. Kubica, Lidia Boghean, Sophie Alvarez, Michael J. Naldrett, Sarbjit Singh, Sandeep Rana, Muhammad Zahid, Janet Schaefer-Klein, Anja Roden, Farhad Kosari, Panos Z. Anastasiadis, Mitesh Borad, Amarnath Natarajan, Aaron S. Mansfield

## Abstract

Despite ongoing efforts to employ structure-based methods to discover targeted protein degraders (TPD), the prevailing strategy continues to be the synthesis of a focused set of heterobifunctional compounds and screen them for target protein degradation. Here we used a fluorescence based live cell imaging screen to identify degraders that target exon 14 skipped hepatocyte growth factor receptor (MET). MET is a known oncogenic driver. MET exon 14 skipping mutations (METex14Δ) are found in lung cancers and result in the loss of a degron that is required for E3-ligase recognition and subsequent ubiquitination, prolonging the half-life and oncogenicity of MET. Since proteolysis targeting chimeras (PROTACs) are heterobifunctional molecules that promote target degradation by the proteosome, we sought to restore degradation of MET lost with METex14Δ using a MET-targeting PROTAC. We generated a library of sixty PROTACs of which 37 used the MET inhibitor capmatinib as the protein of interest targeting ligand. We screened this PROTAC library for targeted degradation of METex14Δ-GFP using live cell imaging. We benchmarked out MET-targeting PROTACs to that of a previously reported MET-targeting PROTAC, SJF8240. Curve fitting live cell imaging data affords determination of time required to degrade 50% of the target protein (DT50), which was used in determining structure activity relationships. A promising candidate, 48-284, identified from the screen, exhibited classic PROTAC characteristics, was > 15-fold more potent than SJF8240, had fewer off targets compared to SJF8240, and degraded MET in multiple cell lines.

## Introduction

Hepatocyte growth factor receptor, more commonly referred to as MET, is a known oncogenic driver in multiple malignancies^1^. Recently the tyrosine kinase inhibitors capmatinib and tepotinib have been approved by the United States Food and Drug Administration (FDA) for the treatment of patients with non-small cell lung cancers that harbor *MET* exon 14 skipping mutations (*MET*ex14Δ)^2, 3^. Mutations that affect the donor or acceptor splice sites of *MET* exon 14 pre-mRNA can lead to skipping of exon 14 during splicing and an mRNA product where exons 13 and 15 are fused^4-6^. Subsequent translation results in a shortened MET protein without its juxtamembrane domain. This juxtamembrane domain includes a degron that is recognized by the E3 ubiquitin-protein ligase casitas B-lineage lymphoma (Cbl). In the absence of the MET degron recognized by Cbl, MET is not readily ubiquitinated, thus prolonging its half-life by avoiding degradation by the proteasome. Protein degradation with proteolysis targeting chimeras (PROTACs) is a viable strategy to degrade oncogenic targets^7, 8^. PROTACs are heterobifunctional molecules that link two protein-binding molecules: one that recognizes a target, and the other that recruits an E3 ubiquitin-protein ligase. Thus, PROTACs can recruit neo substrates to E3 ubiquitin-protein ligases for ubiquitination. Since *MET*ex14Δ results in the loss of the degron that is required for recognition by Cbl and subsequent degradation, we hypothesized that a MET targeting PROTAC could restore degradation of MET with exon 14 skipping mutations. To that end, we synthesized a library of sixty PROTACs of which 37 used tyrosine kinase inhibitor capmatinib. The library was screened using live cell high throughput imaging with *MET*ex14Δ-GFP expressing cell line. This strategy not only allows rapid screening of PROTAC library but also affords structure activity relationship studies using degradation time 50 (DT_50_) derived through curve fitting the time course data. Our studies identified, a promising PROTAC (48-284) that exhibits classical PROTAC phenotype which was more potent and selective compared to the previously reported MET targeted PROTAC SJF8240, and degraded MET in an *in vivo* xenograft model demonstrating target engagement.

## Results

### Synthesis and screening of a PROTAC library to identify MET-Exon14 skipping mutant degraders

To develop MET-targeting PROTACs the tyrosine kinase inhibitor capmatinib was selected, as it had a lower molecular weight, logP, polar surface area and greater potency than tepotinib^9, 10^. Analysis of the Schrödinger GLIDE docked capmatinib into MET (Figure 1A) and the co-crystal structure of a close analog (PDB: 3ZBX)^11^ indicated that the N-methyl amide is solvent exposed and was chosen as the optimal exit vector to introduce the linker. Since linker length and composition play an important role in PROTAC performance, we conjugated the capmatinib acid to an array of linkers through amide chemistry^12, 13^. Thalidomide, lenalidomide and VHL binders were used as E3-targeting ligands to generate a focused set of 37 capmatinib based PROTACs (Scheme S1 and S2). A previously reported MET PROTAC (SJF8240) developed using a promiscuous kinase inhibitor, foretinib^7, 14^ was used as a control. To assess fidelity of the screen we included an additional 23 non-capmatinib based PROTACs with aminopyrazole, palbociclib, futibatinib, BMS345541, YK-4-279 and APS-2-79 as targeting ligands to generate a library of 60 PROTACs (Table S1)^15-18^.

**Fig. 1.**
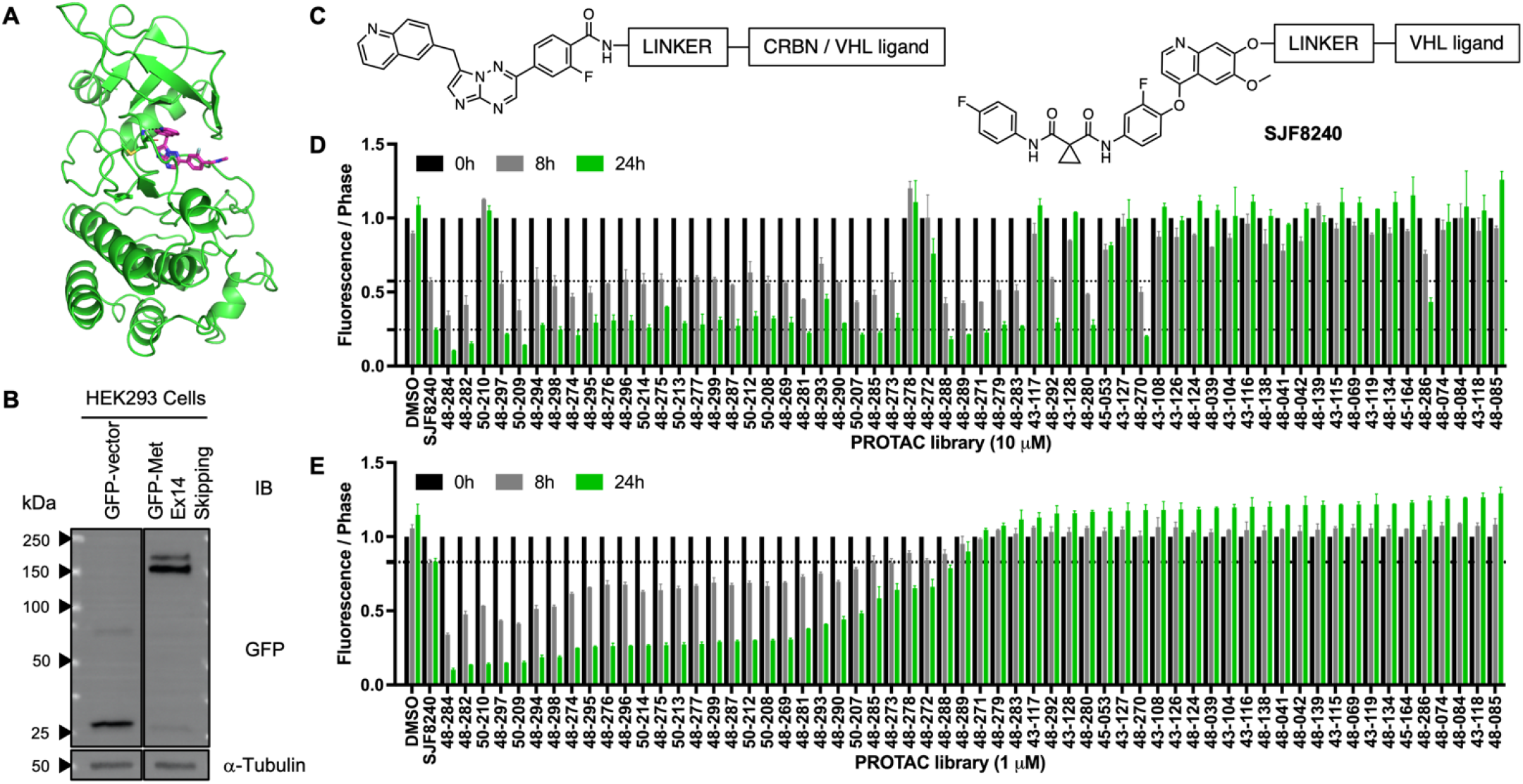
Live cell imaging screen for METex14Δ-GFP degradation. (A) Capmatinib docked into MET kinase domain (pdb: 3zbx). Capmatinib is shown in magenta sticks and the kinase domain of MET is shown as green ribbon structure. The quinoline ring of Capmatinib mimics the adenine ring of the ATP and the quinoline nitrogen is within hydrogen bonding distance of the N-H of Met^1160^ (hydrogen bond is shown in black line) (B) Characterization of HEK293 cells stably transfected with either GFP-vector or GFP-labelled MET-exon-14 skipping mutant. (C) Design of capmatinib based PROTAC library and SJF8240. (D-E) Capmatinib based PROTAC library screened at 10 µM and 1 µM using live cell imaging in HEK293 transfected with GFP-Met-Exon-14 skipping mutant. The bar graph shows the green count values over confluence (phase) normalized to time zero of each well at 0h, 8h and 24h post addition. The broken blackline indicates activity relative to SJF8240. The bars represent mean ± SD of three independent biological replicates (n = 3)

### Live cell imaging-based screen using *MET*ex14Δ-GFP expressing cell line

To rapidly screen the PROTAC library for *MET*ex14Δ degradation, we generated cells that expressed METex14Δ-GFP (Figure 1B). The *MET*ex14Δ-GFP HEK293 cells were used to screen the PROTAC library along with SJF8240 as the positive control at 10, 1 and 0.1 µM in a live cell imaging study (Figure 1C-E, and Figure S1). The treated cells were imaged every 2 hours for GFP signal and confluence (phase)(Videos 1-2). MET-degraders induced a time-dependent loss of GFP signal with minimal effect on the confluence of the cells. The 23 non-capmatinib based PROTACs did not reduce the GFP signal even at 10µM (Figure 1D). Among the capmatinib-based PROTACs, 30/37 exhibited better MET degradation activity at 1µM when compared to SJF8240 (Figure 1E). Rapid degradation of the target protein is a desired feature in PROTACs therefore we determined the time required to degrade 50% of *MET*ex14Δ-GFP (DT_50_) by curve fitting the 1µM time course data (Table S1 and Figure 2A). Based on the DT_50_ values we binned the capmatinib-based PROTACs into 4 groups *viz*., rapid (< 6h), moderate (6-12h), slow (12-22h), and inactive (> 22h).

**Fig. 2.**
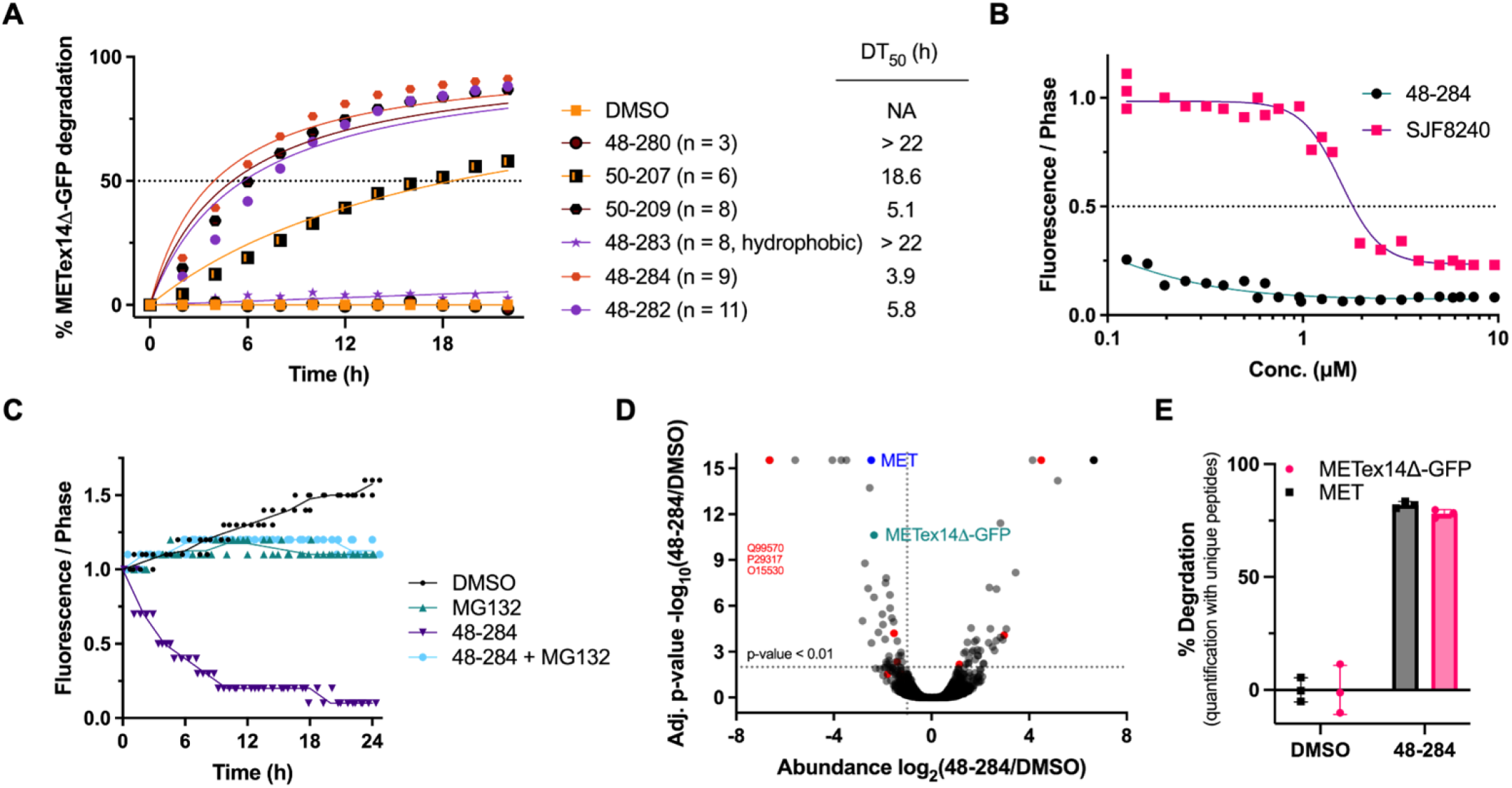
Characterization of MET-PROTACs. (A) Time course study to assess the effects of the linkers in PROTACs generated with capmatinib and VHL ligand. The line graph is an average of three independent biological replicates (n = 3), green count values over confluence (phase) normalized to 0 time of each well. The number of linker atoms are indicated in parathesis. Degradation time for 50% reduction in GFP signal was determined through curve-fitting the data (Prism 10.2.1). (B) A dose-response study with capmatinib based PROTAC 48-284 and foretinib based PROTAC SJF8240 (10000, 5000, 1000, 500, 100 nM) in HEK293 transfected with GFP-Met-Exon-14 skipping mutant (n = 3), green count values over confluence (phase) normalized to time zero of each well. (C) A time course study with the capmatinib based PROTAC 48-284 (1 µM) with HEK293 transfected with GFP-Met-Exon-14 skipping mutant in the presence and absence of MG132 (10 µM). The images analyzed every 2h for 24h post-treatment. The line graph is an average of three independent biological replicates (n = 3), green count values over confluence (phase) normalized to 0 time of each well. (D) Volcano plot depicting changes of protein abundance in HEK293 transfected with GFP-labelled MET-exon-14 skipping mutant cells treated with the PROTAC 48-284 and incubated for 24 hours. The lysates were subjected to label-free proteomic analyses and the volcano plot represents 5106 proteins, with the log2 fold change shown on the x-axis and negative log10 p-values on the y axis. Data are presented as the averages of three independent biological replicates (n = 3). Relevant proteins are labelled. (E) The percent degradation of MET-GFP and GFP-labelled MET-exon-14 skipping is shown for DMSO, and 48-284 with the mean of three replicates shown.

Remarkably, 3 (50-209, 48-284 and 48-282) out of the 4 (48-297) PROTACs that rapidly degraded *MET*ex14Δ-GFP had an all-carbon linker that was conjugated to a VHL ligand *via* an amide bond, while the fourth had a PEG linker conjugated to pomalidomide (Table S1). Among the all-carbon linker VHL ligand PROTACs the 9-atom linker (48-284) was the most potent with a DT_50_ of ∼3.9h. Increasing the linker length to 11-atoms (48-282) or decreasing the linker length to 8-atoms (50-209) resulted in higher DT_50_ values. Irrespective of the length of the linker the PEG-linked VHL ligand PROTACs (48-271, 48-270, 48-283 and 48-286) were all inactive. On the other hand, PROTACs with the PEG linker conjugated to pomalidomide (E1) exhibited an inverse relationship between the DT_50_ values and the linker length with the 14-atom linker (48-297) having the lowest DT_50_ value (5.2h). While the all-carbon linked E1 ligand PROTACs (48-296, 48-299, 48-269, 48-295, and 48-275) exhibited moderate DT_50_ values with no trends. Similarly, we observed no specific trends with PROTACs containing the ether-linked thalidomide. Switching the E3-ligand from pomalidomide (48-298) to lenalidomide (50-214) to ether-linked thalidomide (48-288) while maintaining the linker length and composition resulted in progressive loss of activity (6.866, 9.690, and >22). This suggests that both linker and the E3-ligase ligands play a part in the activity of these capmatinib-based *MET*ex14Δ targeted PROTACs. Based on the activities from the above screen we subjected 48-284, 48-282, and SJF8240, to follow up dose-response and time-course studies (Figure 2B and Figure S2). Consistent with the screening data, 48-284 (DC_50_ < 0.09 μM) was > 15-fold more potent than SJF8240 (DC_50_ = 1.57 μM) (Figure 2B).

To assess if the degradation of *MET*ex14Δ-GFP induced by 48-284 is mediated by the proteasome, METex14Δ-GFP expressing cells were treated with 48-284 in the presence and absence of the proteasome inhibitor MG132. The loss of GFP signal induced by 48-284 was blocked by MG132, suggesting that 48-284 induces proteasomal degradation of METex14Δ-GFP (Figure 2C). To confirm degradation of METex14Δ-GFP and assess the selectivity of 48-284, we subjected METex14Δ-GFP expressing cells to 1 µM of 48-284 for 24h. The lysates from these samples were subjected to mass spectrometry-based proteomic analyses. Conjugating the VHL ligand to capmatinib in 48-284 resulted in 45 proteins identified as hits (abundance > 2-fold reduction and p-value < 0.01). Under the criteria described above, among the 119 kinases quantified, only 4 kinases *viz*., MET, PDPK1, EPHA2 and PIK3R4, were identified as hits in the 48-284 treated samples (Figure 2D). In a similar study with foretinib (a non-selective kinase inhibitor) based MET degrader SJF-8240, degradation of 9 kinases was observed ^14^. Quantification of unique peptides associated with MET and METex14Δ-GFP in samples treated with 48-284 showed that 48-284 potently degraded both MET and METex14Δ-GFP (Figure 2E).

### Validation of 48-284 as a METex14Δ degrader in native METex14Δ expressing cell lines

To confirm the results obtained with 48-284 in the METex14Δ-GFP models, we used cell lines with native *MET*ex14Δ mutations and *MET* amplification. We first selected the Hs746T cell line that was derived from a gastric cancer since this model contains an amplified *MET*ex14Δ mutation^19^. Treatment of Hs746T cells showed a time-dependent decrease in MET when treated with 48-284 at 1.0 µM as assessed by western blot (Figure 3A). There was also a dose-dependent decrease in the levels of MET protein as assessed by western blot, except at levels above 1.0 µM consistent with a hook effect (Figure 3A), a classical feature of a PROTAC. Similar results were obtained with the cell line H596 with a native *MET*ex14Δ mutation (Figure S3).

**Fig. 3.**
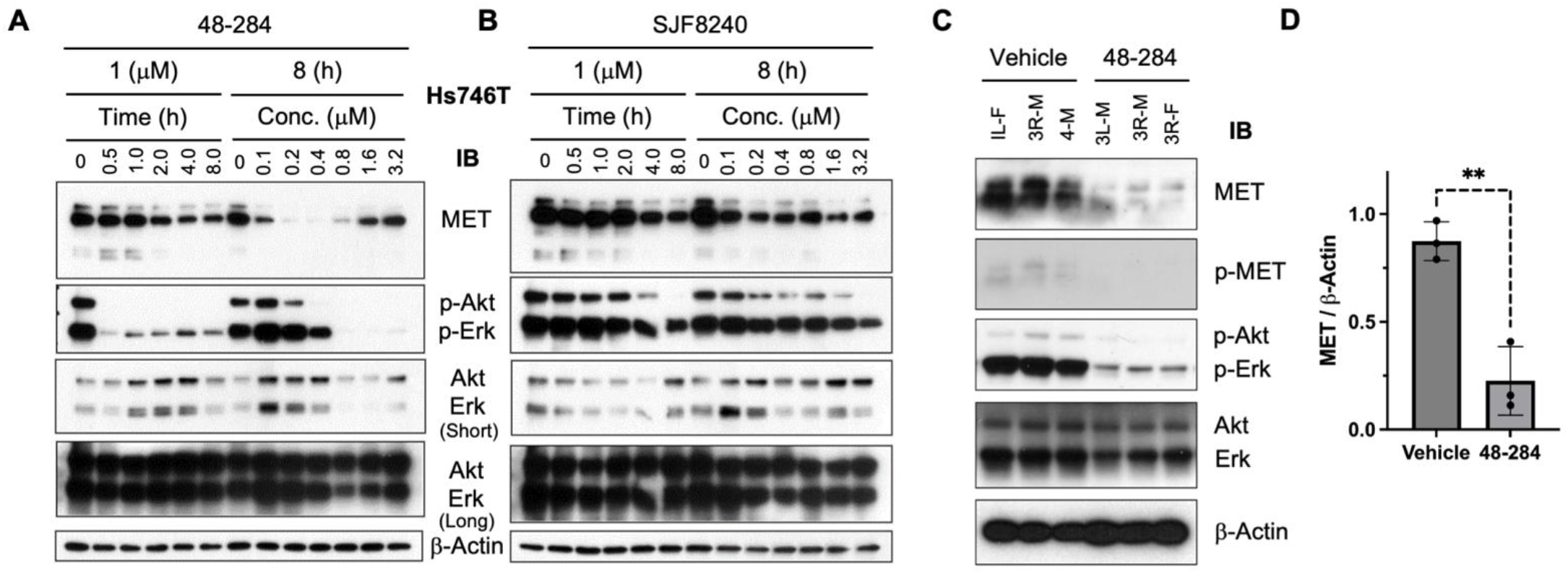
Time and dose effects of MET PROTACs. (A) Hs746T cell line that carries an endogenous *MET*ex14D mutation was treated with capmatinib based PROTAC 48-284 at 1.0 µM and MET was assessed at the indicated timepoints by Western blots with β-actin controls. These cells were also treated at the indicated doses for 8 hours. The effects on downstream RAS/AKT and RAS/ERK pathway signaling were also assessed. (B) Similarly, Hs746T cell line that carries an endogenous *MET*ex14D mutation was treated with the foretinib-based PROTAC SJF-8240 at 1.0 µM and MET was assessed at the indicated timepoints by Western blots with β-actin controls. These cells were also treated at the indicated doses for 8 hours. The effects on downstream RAS/AKT and RAS/ERK pathway signaling were also assessed. (C) Western blot analysis 3 untreated UW21 xenografts (Ctrl) and 3 UW21 xenografts treated with 48-284 (T). We determined the ratios of MET to Beta-actin using ImageJ and found that there was significantly greater MET in the untreated group than treated group (**p-value < 0.01).

Previous work has shown that *MET*ex14Δ increases and prolongs RAS/AKT and RAS/ERK pathway signalling^5^. For that reason, we assessed the effects of 48-284 and SJF8240 on these pathways. We observed reduction of phosphorylation of AKT and MAPK in 48-284 treated cells when compared to SJF-8240 (Figure 3A-B). Together, these studies suggest that 48-284 is a potent degrader of MET, and disrupts MET phosphorylation in cell lines with native *MET*ex14Δ mutations. Consistent with our results in *MET*ex14Δ-GFP cells 48-284 was a more potent degrader when compared to SJF8240. Together these studies validate the use of *MET*ex14Δ-GFP HEK293 cells as a viable screening tool to both identify hit degraders and derive structure activity relationships.

### Validation of target engagement / degradation *in vivo*

To assess target engagement *in vivo* by 48-284, we performed a short perturbation study. We implanted *MET*ex14Δ-mutant UW21 xenografts^20^ into the flanks of mice and allowed them to grow for 2-weeks. Half of the mice were then treated with 48-284 through a tail vein injection twice, eight hours apart, and the tumours were removed six hours after the second injection. Western blot analyses with these tumours showed a decrease in MET in the treated models compared to the untreated models (Figure 3C-D). Consistently, in an immunohistochemistry study, compared to the untreated controls (Figure S4A), the treated tumours (Figure S4B) showed significant reduction in MET expression (p < 0.001, mean percentage of MET in control group 48% versus mean percentage in treated group 22%, Figure S4C).

## Discussion

Targeted protein degradation is emerging as a potent therapeutic tool. The possibilities to degrade undruggable targets and overcome the limitations of occupancy-based efficacy of kinase inhibitors with event-based degradation could potentially change the therapeutic landscape in oncology and other disciplines^21^. Live cell imaging with a fluorescence read out offers a rapid method to not only identify hits but also exploit the time to degrade 50% of the target protein (DT_50_) as a useful tool to conduct structure activity relationship analyses.

MET has been an elusive target for decades despite its known roles across malignancies. It has only been in the last few years that potent, specific MET kinase inhibitors have received accelerated FDA-approval, albeit based on nonrandomized clinical trials. Whereas *MET* amplification or MET expression have not yet clearly translated into predictive biomarkers, *MET*ex14Δ mutations have emerged as a predictive biomarker with MET kinase inhibitors in non-small cell lung cancer. Since the oncogenicity of *MET*ex14Δ mutations is driven by loss of ubiquitination and degradation, we sought to restore the degradation of MET with a PROTAC.

We synthesized a library of MET-targeting PROTACs based on the MET kinase inhibitor capmatinib. In general, the PROTACs with longer hydrophobic carbon linkers had greater activity than shorter linkers. Unlike the foretinib-based MET-targeting PROTAC, inclusion of oxygen atoms in the linkers of capmatinib based MET-targeting PROTACs completely abolished their activity. The degradation of MET that was observed in fluorescent METex14Δ-GFP HEK293T cells was confirmed by mass spectrometry; however, a few other proteins were also degraded. Prior work demonstrated converting kinase inhibitors to PROTACs improves degradation selectivity ^14,15,22^, here we show that using a more selective targeting ligand (capmatinib vs. foretinib) also improves the selectivity profile. Compared to the 9 kinases degraded by SJF8240, only 4 kinases were degraded by 48-284. Of these 4 degraded by 48-284, PDPK1 was recently shown to associate with MET to facilitate the phosphorylation of Akt, and concurrent degradation of MET and PDPK1 would also prevent Src-PDPK1 mediated Akt activation^23^. Consistent with other reported PROTACs^15^, 48-284, also demonstrated the hook-effect at higher concentrations likely due to saturation of the target and E3 ligase, separately. Furthermore, inhibition of the proteasome blocked degradation of MET.

A screening system is useful only if the hits identified, from the screen that used fusion proteins, are validated in cell lines that express native mutant proteins. Unlike the hit KRAS^G12C^ PROTAC that was identified using a KRAS^G12C^-GFP expressing cell line that failed to degrade endogenous KRAS^G12C 24^, we observed degradation of MET in cell lines with native *MET*ex14Δ mutations. We also demonstrated in vivo target engagement and degradation using a human *MET*ex14Δ tumour xenograft model. Overall, these data suggest that MET-targeting PROTACs can be identified by live cell high-throughput imaging using METex14Δ-GFP expressing cell lines. PROTAC 48-284 can restore the degradation of MET that is lost with *MET*ex14Δ mutations.

## Conclusions

Here we report the use of a fluorescence based live cell imaging to screen TPDs. The *MET*ex14Δ-GFP expressing cell line can be used to not only identify hits but also to conduct structure activity relationship studies by leveraging the DT_50_ values. A *MET*ex14Δ targeted PROTAC (48-284) identified using the above screen efficiently degraded the target in cancer cell lines and an *in vivo* tumour model that express native *MET*ex14Δ mutations.

## Supporting information

Control_DMSO

Experimental_48_284

## Author Contribution

J.R. Mallareddy - Conceptualization, methodology, data curation, formal analysis, writing– original draft, review and editing

Lin Yang – Conceptualization, methodology, data curation, formal analysis, writing– original draft, review and editing

Wan-Hsin Lin - Methodology, data curation, formal analysis, writing–review and editing

R. Feathers - Methodology, data curation, formal analysis, writing–review and editing

Jennifer Ayers-Ringler - Methodology, data curation, writing–review and editing

Ezequiel Tolosa - Methodology, data curation, writing–review and editing

Amritha G. Kizhake - Methodology, data curation, writing–review and editing

Smitha Kizhake - Methodology, data curation, writing–review and editing

Sydney P. Kubica - Methodology, data curation, writing–review and editing

Lidia Boghean - Methodology, data curation, writing–review and editing

Sophie Alvarez - Methodology, data curation, writing–review and editing

Michael Naldrett - Methodology, data curation, writing–review and editing Sarbjit

Singh - Methodology, data curation, writing–review and editing

Sandeep Rana - Methodology, data curation, writing–review and editing

Muhammad Zahid - Methodology, data curation, writing–review and editing

Janet Schaefer-Klein - Resources, supervision, project administration, writing–review and editing.

Anja Roden - Methodology, data curation, formal analysis, writing–review and editing

Farhad Kosari- Methodology, data curation, formal analysis, writing–review and editing

Panos Z. Anastasiadis - Methodology, data curation, formal analysis, supervision, writing–review and editing

Mitesh Borad - Conceptualization, resources, supervision, writing–review and editing.

Amarnath Natarajan - Conceptualization, resources, data curation, software, formal analysis, supervision, funding acquisition, methodology, project administration, writing–original draft, review and editing.

Aaron S. Mansfield - Conceptualization, resources, data curation, software, formal analysis, visualization, supervision, funding acquisition, methodology, project administration, writing–original draft, review and editing.

## Conflicts of interest

J.R.M., L. Y., M. B., A.N., and A.S.M. are listed as inventors on PCT Int. Appl. (2023), WO 2023249994 A1

## Data availability

The data supporting this article have been included as part of the Supplementary Information.

## Acknowledgements

This work was supported by the Hallick Family, MET Crusaders, the Barry Family and Mayo Clinic’s Center for Individualized Medicine’s High-Definition Therapeutics and Precision Cancer Therapeutics Programs. The Proteomics & Metabolomics Facility (RRID:SCR_021314), Nebraska Center for Biotechnology at the University of Nebraska-Lincoln, and instrumentation are supported by the Nebraska Research Initiative. The work was also supported in part by University of Nebraska collaborative initiative TSG26784 and NIH grants CA197999, CA251151, CA260749, GM121316, and CA036727. L.B. is supported by T32CA009476. P.Z.A. is supported by NIH R01NS101721. ASM is supported by NCI R21 CA251923, Department of Defence W81XWH-22-1-0021 Concept Award, and a Mark Foundation ASPIRE Award. The authors are thankful to Dr. Baschnagel, Dr. Doebele and Dr. Rudin for sharing xenografts or cell lines for this project. The authors are also grateful to Bobbi Jebens, Sarah A. Anderson, Mark Heggestad and Leila Jones for their administrative support.

## References

1. Guo, R.; Luo, J.; Chang, J.; Rekhtman, N.; Arcila, M.; Drilon, A. MET-dependent solid tumours - molecular diagnosis and targeted therapy. Nat Rev Clin Oncol 2020, 17, 569–587.

2. Paik, P. K.; Felip, E.; Veillon, R.; Sakai, H.; Cortot, A. B.; Garassino, M. C.; Mazieres, J.; Viteri, S.; Senellart, H.; Van Meerbeeck, J.; Raskin, J.; Reinmuth, N.; Conte, P.; Kowalski, D.; Cho, B. C.; Patel, J. D.; Horn, L.; Griesinger, F.; Han, J. Y.; Kim, Y. C.; Chang, G. C.; Tsai, C. L.; Yang, J. C.; Chen, Y. M.; Smit, E. F.; van der Wekken, A. J.; Kato, T.; Juraeva, D.; Stroh, C.; Bruns, R.; Straub, J.; Johne, A.; Scheele, J.; Heymach, J. V.; Le, X. Tepotinib in Non-Small-Cell Lung Cancer with MET Exon 14 Skipping Mutations. N Engl J Med 2020, 383, 931–943.

3. Wolf, J.; Seto, T.; Han, J. Y.; Reguart, N.; Garon, E. B.; Groen, H. J. M.; Tan, D. S. W.; Hida, T.; de Jonge, M.; Orlov, S. V.; Smit, E. F.; Souquet, P. J.; Vansteenkiste, J.; Hochmair, M.; Felip, E.; Nishio, M.; Thomas, M.; Ohashi, K.; Toyozawa, R.; Overbeck, T. R.; de Marinis, F.; Kim, T. M.; Laack, E.; Robeva, A.; Le Mouhaer, S.; Waldron-Lynch, M.; Sankaran, B.; Balbin, O. A.; Cui, X.; Giovannini, M.; Akimov, M.; Heist, R. S.; Investigators, G. m.-. Capmatinib in MET Exon 14-Mutated or MET-Amplified Non-Small-Cell Lung Cancer. N Engl J Med 2020, 383, 944–957.

4. Kong-Beltran, M.; Seshagiri, S.; Zha, J.; Zhu, W.; Bhawe, K.; Mendoza, N.; Holcomb, T.; Pujara, K.; Stinson, J.; Fu, L.; Severin, C.; Rangell, L.; Schwall, R.; Amler, L.; Wickramasinghe, D.; Yauch, R. Somatic mutations lead to an oncogenic deletion of met in lung cancer. Cancer Res 2006, 66, 283–9.

5. Lu, X.; Peled, N.; Greer, J.; Wu, W.; Choi, P.; Berger, A. H.; Wong, S.; Jen, K. Y.; Seo, Y.; Hann, B.; Brooks, A.; Meyerson, M.; Collisson, E. A. MET Exon 14 Mutation Encodes an Actionable Therapeutic Target in Lung Adenocarcinoma. Cancer Res 2017, 77, 4498–4505.

6. Ma, P. C.; Kijima, T.; Maulik, G.; Fox, E. A.; Sattler, M.; Griffin, J. D.; Johnson, B. E.; Salgia, R. c-MET mutational analysis in small cell lung cancer: novel juxtamembrane domain mutations regulating cytoskeletal functions. Cancer Res 2003, 63, 6272–81.

7. Burslem, G. M.; Smith, B. E.; Lai, A. C.; Jaime-Figueroa, S.; McQuaid, D. C.; Bondeson, D. P.; Toure, M.; Dong, H.; Qian, Y.; Wang, J.; Crew, A. P.; Hines, J.; Crews, C. M. The Advantages of Targeted Protein Degradation Over Inhibition: An RTK Case Study. Cell Chem Biol 2018, 25, 67–77 e3.

8. Schapira, M.; Calabrese, M. F.; Bullock, A. N.; Crews, C. M. Targeted protein degradation: expanding the toolbox. Nat Rev Drug Discov 2019, 18, 949–963.

9. Bladt, F.; Faden, B.; Friese-Hamim, M.; Knuehl, C.; Wilm, C.; Fittschen, C.; Gradler, U.; Meyring, M.; Dorsch, D.; Jaehrling, F.; Pehl, U.; Stieber, F.; Schadt, O.; Blaukat, A. EMD 1214063 and EMD 1204831 constitute a new class of potent and highly selective c-Met inhibitors. Clin Cancer Res 2013, 19, 2941–51.

10. Liu, X.; Wang, Q.; Yang, G.; Marando, C.; Koblish, H. K.; Hall, L. M.; Fridman, J. S.; Behshad, E.; Wynn, R.; Li, Y.; Boer, J.; Diamond, S.; He, C.; Xu, M.; Zhuo, J.; Yao, W.; Newton, R. C.; Scherle, P. A. A novel kinase inhibitor, INCB28060, blocks c-MET-dependent signaling, neoplastic activities, and cross-talk with EGFR and HER-3. Clin Cancer Res 2011, 17, 7127–38.

11. Cui, J. J.; Shen, H.; Tran-Dube, M.; Nambu, M.; McTigue, M.; Grodsky, N.; Ryan, K.; Yamazaki, S.; Aguirre, S.; Parker, M.; Li, Q.; Zou, H.; Christensen, J. Lessons from (S)-6-(1-(6-(1-methyl-1H-pyrazol-4-yl)-[1,2,4]triazolo[4,3-b]pyridazin-3-yl)ethyl)quinoline (PF-04254644), an inhibitor of receptor tyrosine kinase c-Met with high protein kinase selectivity but broad phosphodiesterase family inhibition leading to myocardial degeneration in rats. J Med Chem 2013, 56, 6651–65.

12. Churcher, I. Protac-Induced Protein Degradation in Drug Discovery: Breaking the Rules or Just Making New Ones? J Med Chem 2018, 61, 444–452.

13. King, H. M.; Rana, S.; Kubica, S. P.; Mallareddy, J. R.; Kizhake, S.; Ezell, E. L.; Zahid, M.; Naldrett, M. J.; Alvarez, S.; Law, H. C.; Woods, N. T.; Natarajan, A. Aminopyrazole based CDK9 PROTAC sensitizes pancreatic cancer cells to venetoclax. Bioorg Med Chem Lett 2021, 43, 128061.

14. Bondeson, D. P.; Smith, B. E.; Burslem, G. M.; Buhimschi, A. D.; Hines, J.; Jaime-Figueroa, S.; Wang, J.; Hamman, B. D.; Ishchenko, A.; Crews, C. M. Lessons in PROTAC Design from Selective Degradation with a Promiscuous Warhead. Cell Chem Biol 2018, 25, 78–87 e5.

15. Rana, S.; Bendjennat, M.; Kour, S.; King, H. M.; Kizhake, S.; Zahid, M.; Natarajan, A. Selective degradation of CDK6 by a palbociclib based PROTAC. Bioorg Med Chem Lett 2019, 29, 1375–1379.

16. Tosso, P. N.; Kong, Y.; Scher, L.; Cummins, R.; Schneider, J.; Rahim, S.; Holman, K. T.; Toretsky, J.; Wang, K.; Uren, A.; Brown, M. L. Synthesis and structure-activity relationship studies of small molecule disruptors of EWS-FLI1 interactions in Ewing’s sarcoma. J Med Chem 2014, 57, 10290–303.

17. Dhawan, N. S.; Scopton, A. P.; Dar, A. C. Small molecule stabilization of the KSR inactive state antagonizes oncogenic Ras signalling. Nature 2016, 537, 112–116.

18. Napoleon, J. V.; Singh, S.; Rana, S.; Bendjennat, M.; Kumar, V.; Kizhake, S.; Palermo, N. Y.; Ouellette, M. M.; Huxford, T.; Natarajan, A. Small molecule binding to inhibitor of nuclear factor kappa-B kinase subunit beta in an ATP non-competitive manner. Chem Commun (Camb) 2021, 57, 4678–4681.

19. Asaoka, Y.; Tada, M.; Ikenoue, T.; Seto, M.; Imai, M.; Miyabayashi, K.; Yamamoto, K.; Yamamoto, S.; Kudo, Y.; Mohri, D.; Isomura, Y.; Ijichi, H.; Tateishi, K.; Kanai, F.; Ogawa, S.; Omata, M.; Koike, K. Gastric cancer cell line Hs746T harbors a splice site mutation of c-Met causing juxtamembrane domain deletion. Biochem Biophys Res Commun 2010, 394, 1042–6.

20. Baschnagel, A. M.; Kaushik, S.; Durmaz, A.; Goldstein, S.; Ong, I. M.; Abel, L.; Clark, P. A.; Gurel, Z.; Leal, T.; Buehler, D.; Iyer, G.; Scott, J. G.; Kimple, R. J. Development and characterization of patient-derived xenografts from non-small cell lung cancer brain metastases. Sci Rep 2021, 11, 2520.

21. Bekes, M.; Langley, D. R.; Crews, C. M. PROTAC targeted protein degraders: the past is prologue. Nat Rev Drug Discov 2022.

22. Robb, C. M.; Contreras, J. I.; Kour, S.; Taylor, M. A.; Abid, M.; Sonawane, Y. A.; Zahid, M.; Murry, D. J.; Natarajan, A.; Rana, S. Chemically induced degradation of CDK9 by a proteolysis targeting chimera (PROTAC). Chem Commun (Camb) 2017, 53, 7577–7580.

23. Molinaro, C.; Martoriati, A.; Lescuyer, A.; Fliniaux, I.; Tulasne, D.; Cailliau, K. 3-phosphoinositide-dependent protein kinase 1 (PDK1) mediates crosstalk between Src and Akt pathways in MET receptor signaling. FEBS Lett 2021, 595, 2655–2664.

24. Zeng, M.; Xiong, Y.; Safaee, N.; Nowak, R. P.; Donovan, K. A.; Yuan, C. J.; Nabet, B.; Gero, T. W.; Feru, F.; Li, L.; Gondi, S.; Ombelets, L. J.; Quan, C.; Janne, P. A.; Kostic, M.; Scott, D. A.; Westover, K. D.; Fischer, E. S.; Gray, N. S. Exploring Targeted Degradation Strategy for Oncogenic KRAS(G12C). Cell Chem Biol 2020, 27, 19–31 e6.

25. Spigel, D. R.; Edelman, M. J.; O’Byrne, K.; Paz-Ares, L.; Mocci, S.; Phan, S.; Shames, D. S.; Smith, D.; Yu, W.; Paton, V. E.; Mok, T. Results From the Phase III Randomized Trial of Onartuzumab Plus Erlotinib Versus Erlotinib in Previously Treated Stage IIIB or IV Non-Small-Cell Lung Cancer: METLung. J Clin Oncol 2017, 35, 412–420.

26. Drilon, A.; Clark, J. W.; Weiss, J.; Ou, S. I.; Camidge, D. R.; Solomon, B. J.; Otterson, G. A.; Villaruz, L. C.; Riely, G. J.; Heist, R. S.; Awad, M. M.; Shapiro, G. I.; Satouchi, M.; Hida, T.; Hayashi, H.; Murphy, D. A.; Wang, S. C.; Li, S.; Usari, T.; Wilner, K. D.; Paik, P. K. Antitumor activity of crizotinib in lung cancers harboring a MET exon 14 alteration. Nat Med 2020, 26, 47–51.

