## Supplemental_Information for "Fluorescence based live cell imaging identifies exon 14 skipped hepatocyte growth factor receptor (MET) degraders"

[a.] *Eppley Institute for Cancer Research, University of Nebraska Medical Center, Omaha NE 68198.*
[b.] *Precision Cancer Therapeutics, Center for Individualized Medicine, Mayo Clinic, Rochester, MN 55905.*
[c.] *Proteomics & Metabolomics Facility, Nebraska Center for Biotechnology, University of Nebraska, Lincoln, NE 68588.*
[d.] *Department of Environmental, Agricultural and Occupational Health, University of Nebraska Medical Center, Omaha NE 68198.*
[e.] *Department of Laboratory Medicine and Pathology, Mayo Clinic, Rochester, MN 55905.*
[f.] *Division of Medical Oncology, Mayo Clinic, Phoenix, AZ 85054, USA*
[g.] *Department of Pharmaceutical Sciences, Department of Genetics Cell Biology & Anatomy, Fred & Pamela Buffett Cancer Center, University of Nebraska Medical Center, Omaha NE 68198*
[h.] *Division of Medical Oncology, Mayo Clinic, Rochester, MN 55905, USA*
### Equal contribution.
Supplementary Information available: [details of any supplementary information available should be included here]. See DOI: 10.1039/x0xx00000x

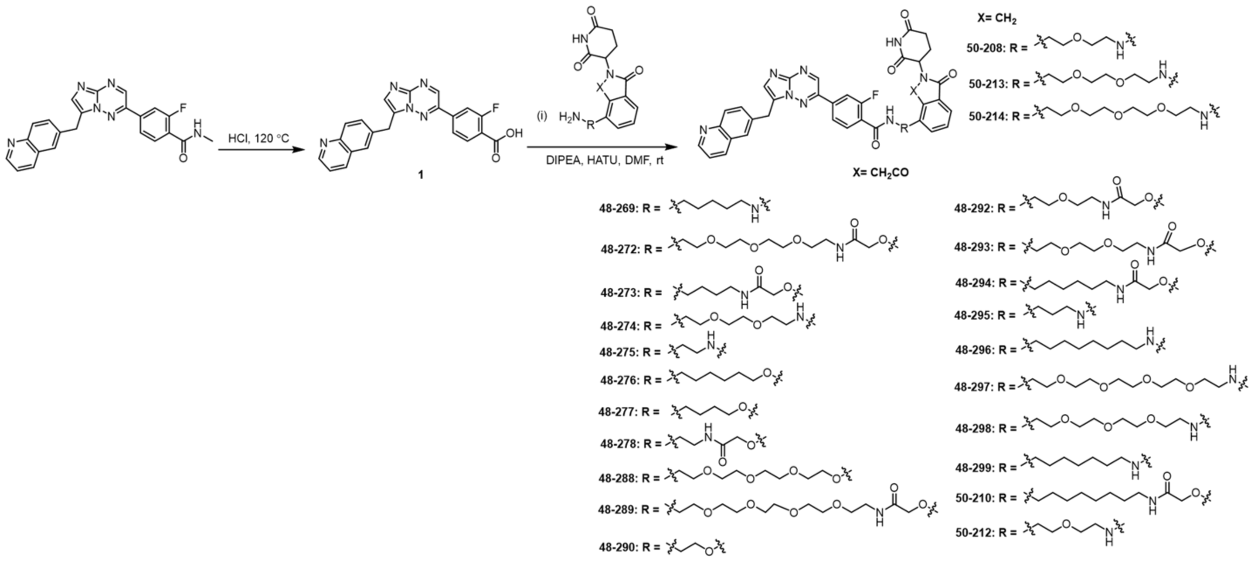

**Scheme S1**. Synthesis of 2-fluoro-4-(7-(quinolin-6-ylmethyl)imidazo[1,2-b][1,2,4]triazin-2-yl)benzoic acid (Capmatinib) based PROTACs with amine-containing linkers conjugated to Thalidomide or Pomalidomide or Lenalidomide ligand

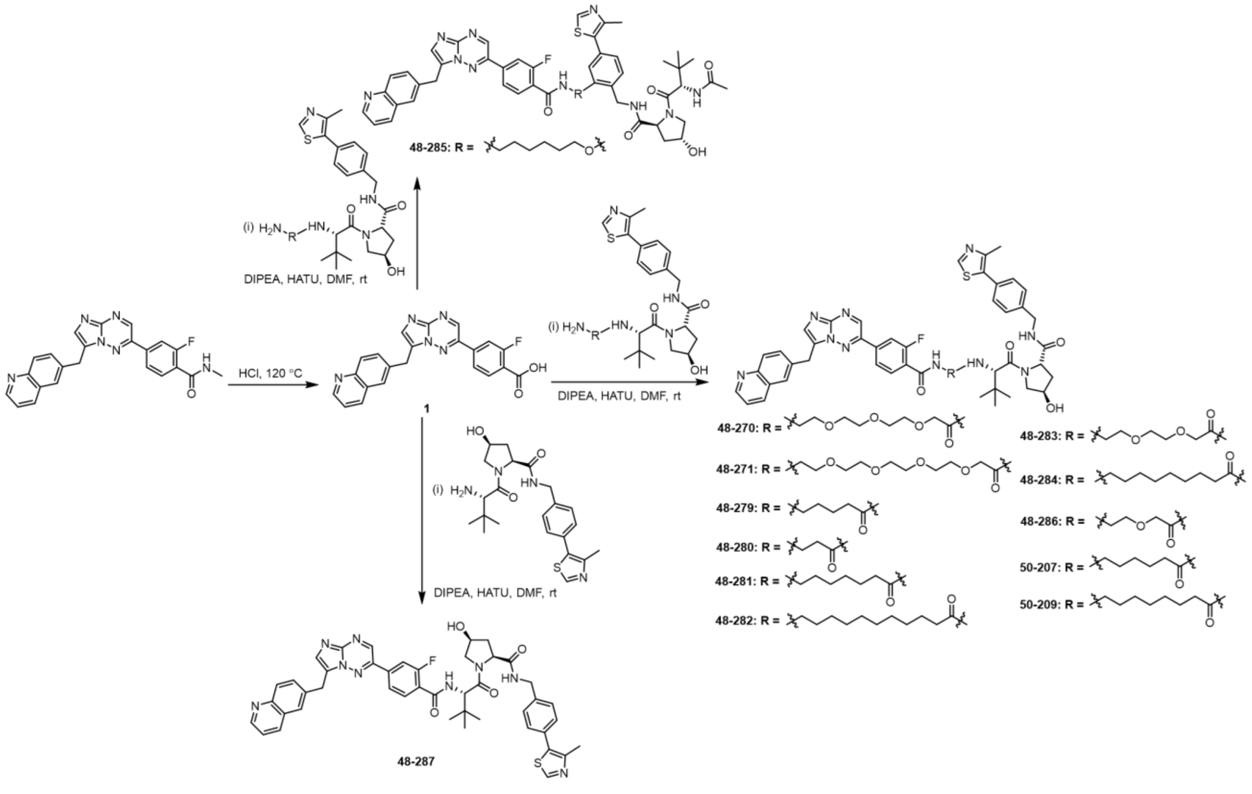

**Scheme S2**. Synthesis of 2-fluoro-4-(7-(quinolin-6-ylmethyl)imidazo[1,2-b][1,2,4]triazin-2-yl)benzoic acid (Capmatinib) based PROTACs with amine-containing linkers conjugated to VHL ligand

**Table S1**. PROTAC library with different head groups (HGs), linkers, and E3 ligands.

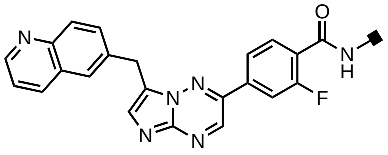

**HG-1**

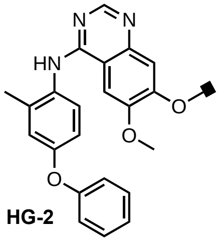

**HG-2**

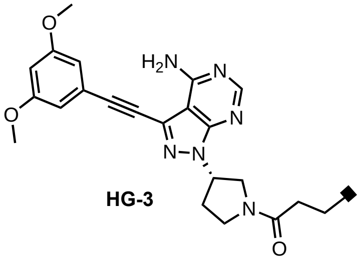

**HG-3**

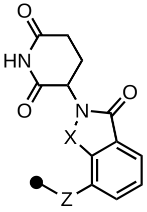

**E-1**; X = CO; Z = NH
**E-2**; X = $CH_2$; Z = NH
**E-3**; X = CO; Z = O

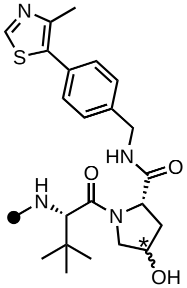

**E-4**; * = S
**E-5**; * = R

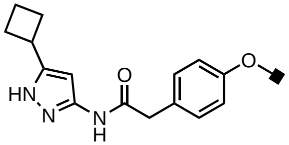

**HG-4**

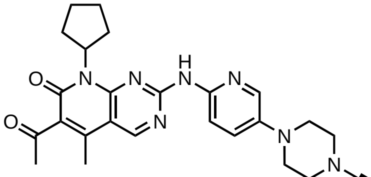

**HG-5**

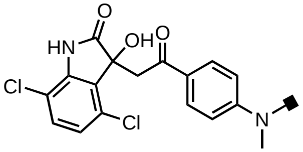

**HG-6**

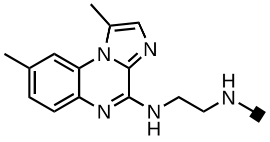

**HG-7**

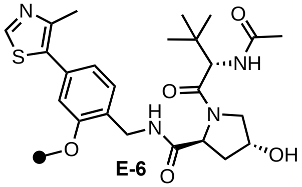

**E-6**

| Composition | Compound | Head Group | E3 Ligand | DT$_{50}$ (h)[#] | MW | Linker |
|---|---|---|---|---|---|---|
| PEG | 48-297 | HG1 | E1 | 5.193 | 874 | 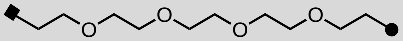 |
| | 48-298 | HG1 | E1 | 6.866 | 830 | 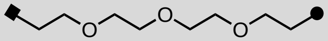 |
| | 48-274 | HG1 | E1 | 9.021 | 786 | 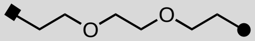 |
| | 50-212 | HG1 | E1 | 11.44 | 742 | 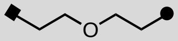 |
| Carbon | 48-296 | HG1 | E1 | 10.68 | 782 | 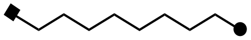 |
| | 48-299 | HG1 | E1 | 11.42 | 768 | 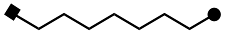 |
| | 48-269 | HG1 | E1 | 11.79 | 740 | 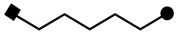 |

| | | | | | | |
|---|---|---|---|---|---|---|
| | 48-295 | HG1 | E1 | 10.23 | 712 | 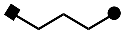 |
| | 48-275 | HG1 | E1 | 9.782 | 698 | 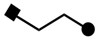 |
| PEG | 50-214 | HG1 | E2 | 9.690 | 816 | 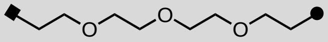 |
| | 50-213 | HG1 | E2 | 10.29 | 772 | 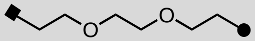 |
| | 50-208 | HG1 | E2 | 11.13 | 728 | 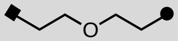 |
| | 48-288 | HG1 | E3 | >22 | 831 | 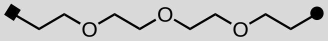 |
| Carbon | 48-276 | HG1 | E3 | 10.53 | 755 | 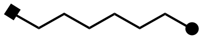 |
| | 48-277 | HG1 | E3 | 10.69 | 727 | 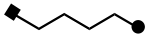 |
| | 48-290 | HG1 | E3 | 14.09 | 699 | 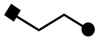 |
| Mixed PEG | 48-289 | HG1 | E3 | >22 | 932 | 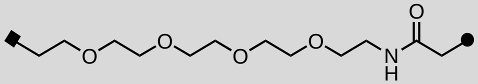 |
| | 48-272 | HG1 | E3 | >22 | 888 | 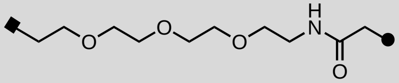 |
| | 48-293 | HG1 | E3 | 15.76 | 844 | 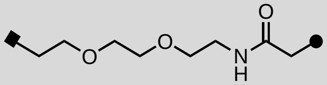 |
| | 48-292 | HG1 | E3 | >22 | 800 | 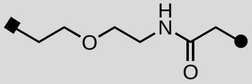 |
| Mixed Carbon | 50-210 | HG1 | E3 | 6.482 | 840 | 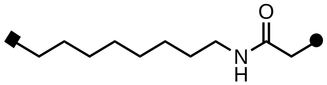 |

| | | | | | | |
|---|---|---|---|---|---|---|
| | 48-294 | HG1 | E3 | 6.547 | 812 | 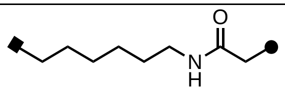 |
| | 48-273 | HG1 | E3 | >22 | 784 | 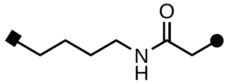 |
| | 48-278 | HG1 | E3 | >22 | 756 | 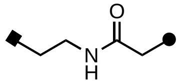 |
| | 48-271 | HG1 | E4 | >22 | 1045 | 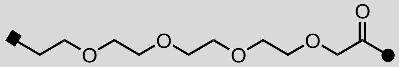 |
| Mixed PEG | 48-270 | HG1 | E4 | >22 | 1001 | 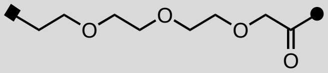 |
| | 48-283 | HG1 | E4 | >22 | 957 | 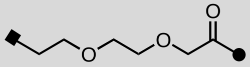 |
| | 48-286 | HG1 | E4 | >22 | 913 | 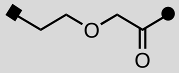 |
| | 48-282 | HG1 | E4 | 5.755 | 995 | 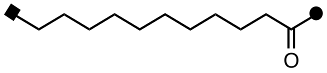 |
| | 48-284 | HG1 | E4 | 3.932 | 967 | 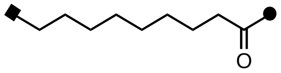 |
| Mixed Carbon | 50-209 | HG1 | E4 | 5.057 | 953 | 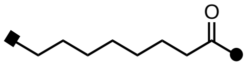 |
| | 48-281 | HG1 | E4 | 14.00 | 939 | 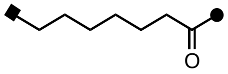 |
| | 50-207 | HG1 | E4 | 18.58 | 925 | 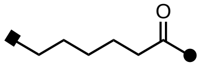 |

| | | | | | | |
|---|---|---|---|---|---|---|
| | 48-279 | HG1 | E4 | >22 | 911 | 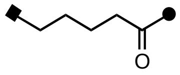 |
| | 48-280 | HG1 | E4 | >22 | 883 | 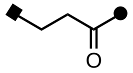 |
| | 48-287* | HG1 | E5 | 11.05 | 812 | |
| Carbon | 48-285 | HG1 | E6 | >22 | 969 | 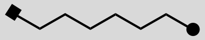 |
| Mixed PEG | 48-041 | HG2 | E1 | >22 | 962 | 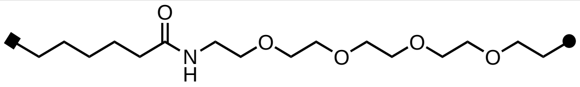 |
| | 48-042 | HG2 | E1 | >22 | 872 | 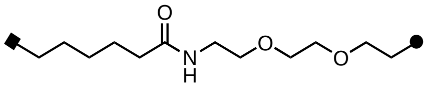 |
| Mixed Carbon | 48-039 | HG2 | E1 | >22 | 828 | 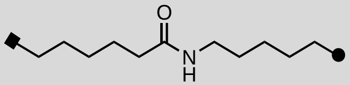 |
| Mixed PEG | 48-138 | HG3 | E1 | >22 | 955 | 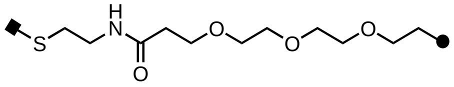 |
| Mixed Carbon | 48-139 | HG3 | E1 | >22 | 865 | 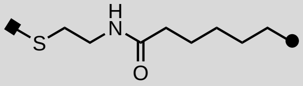 |
| | 48-134 | HG3 | E1 | >22 | 809 | 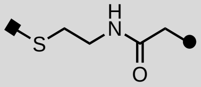 |
| Mixed PEG | 43-128 | HG4 | E1 | >22 | 920 | 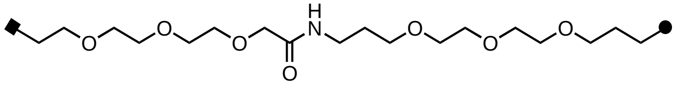 |
| | 43-127 | HG4 | E1 | >22 | 848 | 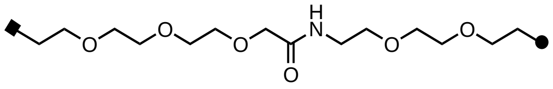 |

| | | | | | | |
|---|---|---|---|---|---|---|
| | 43-119 | HG4 | E1 | >22 | 816 | 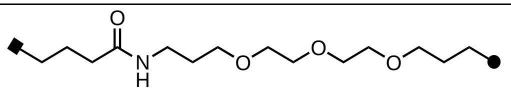 |
| | 43-104 | HG4 | E1 | >22 | 788 | 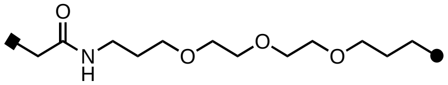 |
| | 43-126 | HG4 | E1 | >22 | 788 | 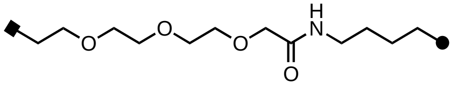 |
| | 43-118 | HG4 | E1 | >22 | 744 | 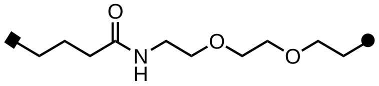 |
| | 43-116 | HG4 | E1 | >22 | 716 | 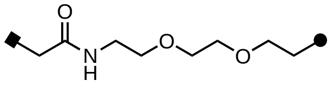 |
| Mixed Carbon | 43-117 | HG4 | E1 | >22 | 684 | 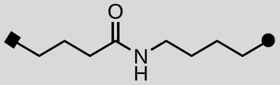 |
| | 43-108 | HG4 | E1 | >22 | 712 | 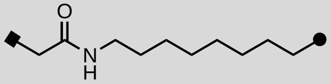 |
| | 43-115 | HG4 | E1 | >22 | 656 | 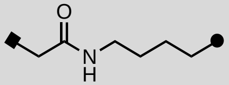 |
| Mixed PEG | 48-124 | HG5 | E4 | >22 | 1151 | 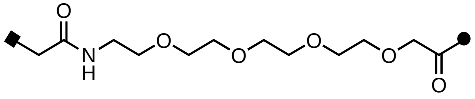 |
| | 45-053 | HG5 | E1 | >22 | 964 | 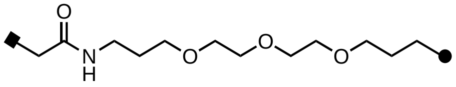 |

| | | | | | | |
|---|---|---|---|---|---|---|
| Mixed Carbon | 45-164 | HG6 | E1 | >22 | 792 | 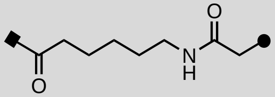 |
| Mixed PEG | 48-074 | HG7 | E1 | >22 | 814 | 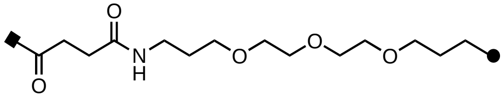 |
| | 48-084 | HG7 | E1 | >22 | 715 | 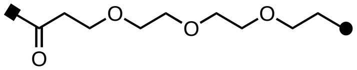 |
| Mixed Carbon | 48-085 | HG7 | E1 | >22 | 625 | 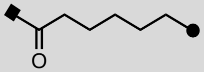 |
| | 48-069 | HG7 | E1 | >22 | 569 | 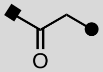 |

[#]Determined by curve fitting the time course data generated by treating *MET*ex14Δ-GFP cells with 1mM of the PROTAC. Head Groups: HG1-capmatinib, HG2- APS-2-79, HG3-futibatinib, HG4-aminopyrazole, HG5-palbociclib, HG6- YK-4-279, HG7-BMS345541. *A direct carboxamide bond between HG1 and E4.

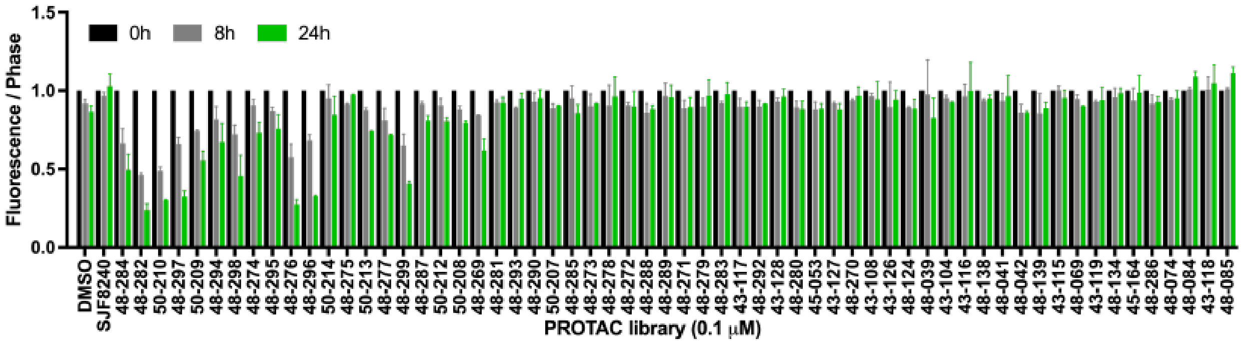


**Figure S1.** Capmatinib based PROTAC library screened at 0.1 μM using live cell imaging in HEK293 transfected with GFP-Met-Exon-14 skipping mutant. The bar graph shows the green count values over confluence (phase) normalized to time zero of each well at 0h, 8h and 24h post addition. The bars represent mean ± SD of three independent biological replicates (n = 3).

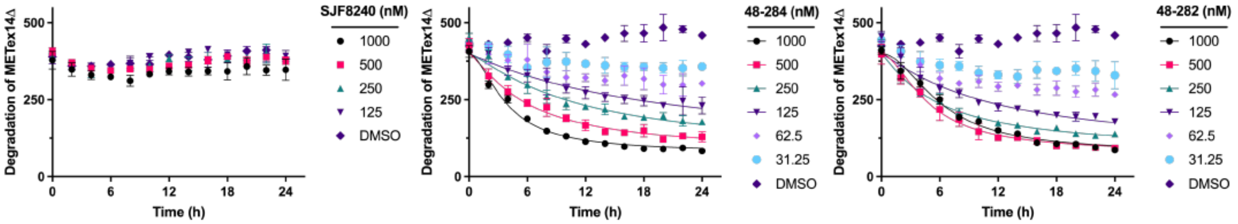


**Figure S2.** A dose-response and time course study using live cell imaging in HEK293 transfected with GFP-Met-Exon-14 skipping mutant treated with SJF8240, 48-284 and 48-282. The line graph shows the green count values over confluence (phase) normalized to time zero. The data is represented mean ± SD of three independent biological replicates (n = 3).

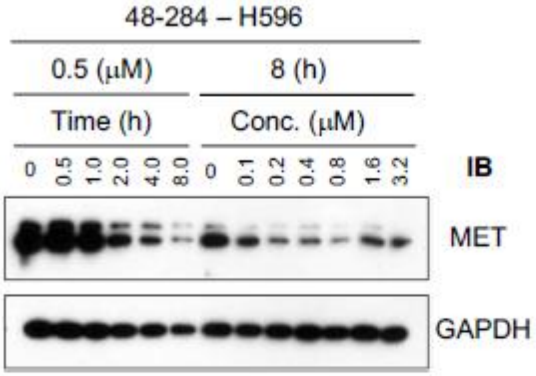


**Figure S3.** Western blot for the cell lines H596 which has a native METex14Δ mutation and was treated with the MET targeting PROTAC 48-284 at 0.5 μM over 8 hours. This cell line was also treated at increasing concentrations for 8 hours as labeled with GAPDH controls.

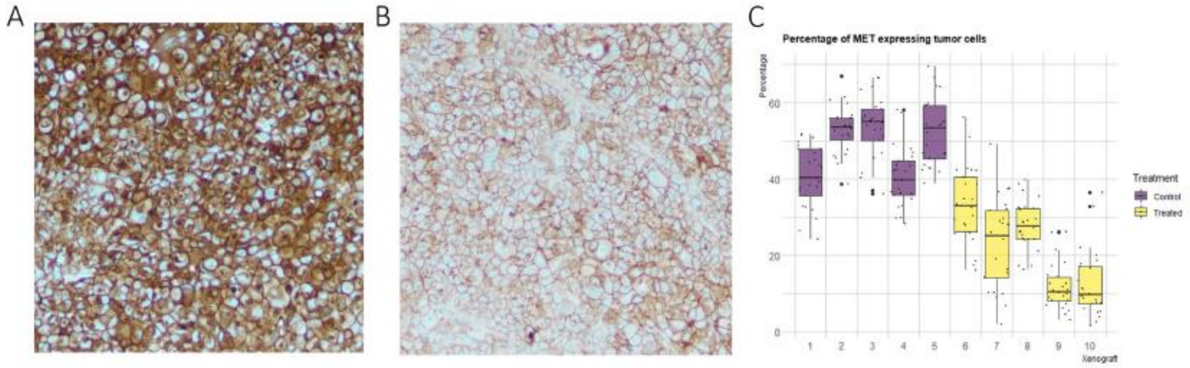


**Figure S4.** (A) Multiple UW21 xenografts with a METex14Δ mutation were removed from mouse flanks and stained for MET by immunohistochemistry. In this representative example, intense membranous and cytoplasmic MET expression is seen in almost all the tumor cells in the control group treated with DMSO. (B) The other group of UW21 xenografts were treated with two doses of 48-284 through tail vein injection eight hours apart, removed six hours after the second injection and stained for MET by immunohistochemistry. In this representative example, significant reduction in MET expression was observed along with evidence of a treatment effect. (C) The percentage of MET expressing tumor cells was quantified digitally at multiple regions of interest in each control and treated xenografts and visualized with this boxplot created with R.

#### MATERIALS AND METHODS

#### General

All reagents were purchased from commercial sources and were used without further purification. $^{1}H$ NMR and $^{13}C$ NMR spectra were recorded in DMSO-$d_6$ on a Varian-500, Varian-600 and Bruker 400 (DMSO-$d_6$ 2.50 ppm for $^{1}H$ and 39.00 ppm for $^{13}C$). Proton and carbon chemical shifts were reported in ppm relative to the signal from residual solvent proton and carbon. The purity of all final compounds was ≥96% as determined by analytical HPLC on a reverse-phase column (Phenomenex C18, 150x4.6 mm, 5 µm) using a Waters 2695 series system with UV detector, l=254nm (Supplemental Information). The eluents: solvent A ($H_2O$ with 0.1% Formic acid) and solvent B ($CH_3CN$ with 0.1% Formic acid), gradient: 10 – 100% B over 15 min with flow rate of 1.0 mL/min. HRMS data was generated on an Agilent 6230 LC/TOF system with UV detector (254 nm).

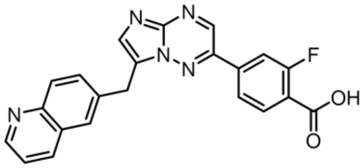


Synthesis of 2-fluoro-4-(7-(quinolin-6-ylmethyl)imidazo[1,2-b][1,2,4]triazin-2-yl)benzoic acid (1)

The 2-fluoro-N-methyl-4-(7-(quinolin-6-ylmethyl)imidazo[1,2-b][1,2,4]triazin-2-yl)benzamide (Capmatinib) (1.0 g) was taken in a round bottom flask, added 37% HCl (8 mL) and heated the reaction mixture at 120°C for 24h. The solution was cooled down to room temperature and slowly added water. To this solution was added 5M KOH slowly until bright fluorescent precipitate was observed. The resulting solution was stirred at 0°C for 20 minutes and precipitate was filtered and air dried to yield product as fluorescent solid. Yield: 730 mg (75%). $^{1}H$ NMR (400 MHz, DMSO-$d_6$) δ 13.54 (s, 1H), 9.24 (s, 1H), 8.95 (dd, $J$ = 4.4, 1.7 Hz, 1H), 8.51 (dd, $J$ = 8.3, 1.6 Hz, 1H), 8.10 – 8.00 (m, 6H), 7.89 (dd, $J$ = 8.6, 2.0 Hz, 1H), 7.64 (dd, $J$ = 8.3, 4.5 Hz, 1H), 4.67 (s, 2H). $^{13}C$ NMR (100 MHz, DMSO-$d_6$) δ 164.97, 164.94, 163.03, 160.47, 149.35, 144.81, 144.16, 141.74, 141.23, 139.01, 138.92, 138.58, 136.94, 135.30, 133.23, 132.56, 128.51, 127.92, 127.65, 126.65, 123.18, 123.15, 122.26, 121.37, 121.26, 115.97, 115.72, 29.16. HRMS (ESI-MS) calcd for $C_{22}H_{15}FN_5O_2^+$ m/z $(M+H)^+$ 400.1204, found: 400.1212.

General procedure for the synthesis of Capmatinib based PROTACs

To a solution of 2-fluoro-4-(7-(quinolin-6-ylmethyl)imidazo[1,2-b][1,2,4]triazin-2-yl)benzoic acid (1.0 eq) and amine that was attached to E3-ligase ligand through linker (1.0 eq) in DMF (1 mL) was added DIPEA (4.0 eq) and stirred for 10 minutes followed by addition of HATU (2.0 eq). The reaction mixture was stirred at room temperature for 16h. The crude was purified by reverse phase HPLC (RP-HPLC) with gradient of 10-100% $CH_3CN$ in $H_2O$ and lyophilized to give product as fluorescent solids. Yields: 40-90%.

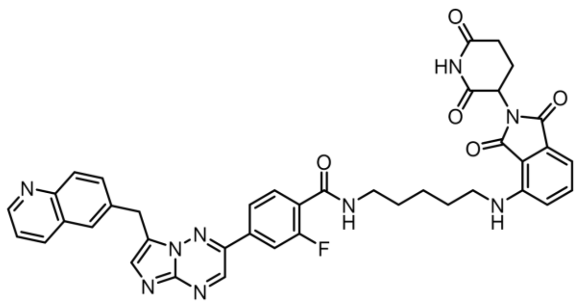


N-(5-((2-(2,6-dioxopiperidin-3-yl)-1,3-dioxoisoindolin-4-yl)amino)pentyl)-2-fluoro-4-(7-(quinolin-6-ylmethyl)imidazo[1,2-b][1,2,4]triazin-2-yl)benzamide (48-269)

Yield = 73%. $^1$H NMR (600 MHz, DMSO-$d_6$) δ 11.08 (s, 1H), 9.18 (s, 1H), 8.97 (dd, *J* = 4.7, 1.6 Hz, 1H), 8.62 (dd, *J* = 8.6, 1.6 Hz, 1H), 8.47 (dt, *J* = 5.9, 3.0 Hz, 1H), 8.11 – 8.03 (m, 2H), 8.02 – 7.90 (m, 4H), 7.76 – 7.69 (m, 2H), 7.56 (dd, *J* = 8.6, 7.1 Hz, 1H), 7.09 (d, *J* = 8.6 Hz, 1H), 7.00 (d, *J* = 7.0 Hz, 1H), 5.00 (dd, *J* = 12.9, 5.5 Hz, 1H), 4.68 (s, 2H), 3.32 – 3.24 (m, 4H), 2.89 – 2.78 (m, 1H), 2.62 – 2.60 (m, 2iH), 1.99 (dtd, *J* = 13.0, 5.5, 2.5 Hz, 1H), 1.59 (dp, *J* = 26.3, 7.2 Hz, 4H), 1.40 (qd, *J* = 9.7, 9.0, 6.2 Hz, 2H). HRMS (ESI-MS) calcd for $C_{40}H_{35}FN_9O_5^+$ m/z $(M+H)^+$ 740.2740, found: 740.2742.

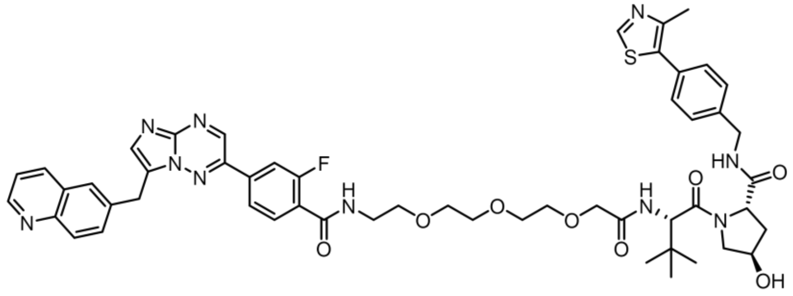


(2S,4R)-1-((S)-15-(tert-butyl)-1-(2-fluoro-4-(7-(quinolin-6-ylmethyl)imidazo[1,2-b][1,2,4]triazin-2-yl)phenyl)-1,13-dioxo-5,8,11-trioxa-2,14-diazahexadecan-16-oyl)-4-hydroxy-N-(4-(4-methylthiazol-5-yl)benzyl)pyrrolidine-2-carboxamide (48-270)

Yield = 60%. $^1$H NMR (600 MHz, DMSO-$d_6$) δ 9.19 (s, 1H), 9.02 (dd, *J* = 4.8, 1.6 Hz, 1H), 8.92 (s, 1H), 8.74 – 8.69 (m, 1H), 8.59 (t, *J* = 6.1 Hz, 1H), 8.44 (qd, *J* = 7.5, 5.9, 2.3 Hz, 1H), 8.14 (d, *J* = 1.8 Hz, 1H), 8.08 (d, *J* = 8.7 Hz, 1H), 8.03 – 7.94 (m, 4H), 7.81 – 7.73 (m, 2H), 7.42 (d, *J* = 9.6 Hz, 1H), 7.40 – 7.32 (m, 4H), 4.68 (s, 2H), 4.53 (d, *J* = 9.6 Hz, 1H), 4.41 (t, *J* = 8.2 Hz, 1H), 4.41 – 4.31 (m, 2H), 4.23 (td, *J* = 15.8, 15.1, 5.5 Hz, 1H), 3.94 (s, 2H), 3.65-3.62 (m, 9), 3.58 – 3.48 (m, 4H), 3.41 (p, *J* = 5.9 Hz, 2H), 2.54 (s, 2H), 2.39 (s, 3H), 2.05 (m 1H), 1.89 (m, 8.9, 4.5 Hz, 1H), 0.90 (s, 9H). HRMS (ESI-MS) calcd for $C_{52}H_{58}FN_{10}O_8S^+$ m/z $(M+H)^+$ 1001.4138, found: 1001.4138.

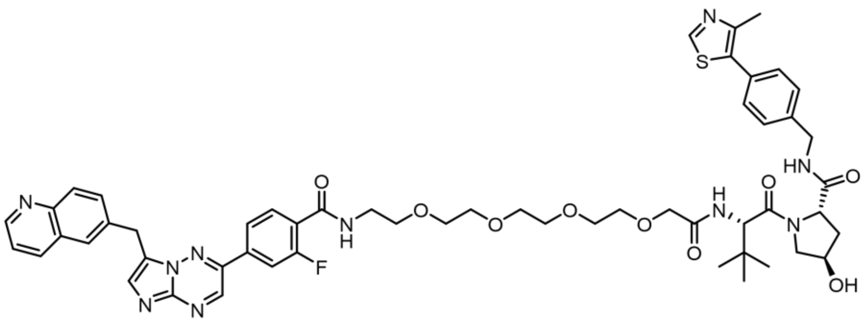

(2S,4R)-1-((S)-18-(tert-butyl)-1-(2-fluoro-4-(7-(quinolin-6-ylmethyl)imidazo[1,2-b][1,2,4]triazin-2-yl)phenyl)-1,16-dioxo-5,8,11,14-tetraoxa-2,17-diazanonadecan-19-oyl)-4-hydroxy-N-(4-(4-methylthiazol-5-yl)benzyl)pyrrolidine-2-carboxamide (48-271)

Yield = 40%. $^1$H NMR (600 MHz, DMSO-$d_6$) δ 9.18 (s, 1H), 9.00 (dd, *J* = 4.8, 1.6 Hz, 1H), 8.93 (s, 1H), 8.69 (dd, *J* = 8.5, 1.5 Hz, 1H), 8.60 (t, *J* = 6.1 Hz, 1H), 8.45 (td, *J* = 5.6, 2.4 Hz, 1H), 8.12 (d, *J* = 1.8 Hz, 1H), 8.07 (d, *J* = 8.7 Hz, 1H), 8.02 – 7.96 (m, 4H), 7.79 – 7.74 (m, 2H), 7.40 (dd, *J* = 8.9, 4.6 Hz, 1H), 7.36 (q, *J* = 8.3 Hz, 4H), 4.68 (s, 2H), 4.53 (d, *J* = 9.6 Hz, 1H), 4.45 – 4.32 (m, 3H), 4.23 (td, *J* = 16.5, 15.8, 5.6 Hz, 1H), 3.93 (s, 2H), 3.57 (dd, *J* = 6.2, 3.4 Hz, 1H), 3.-68-3.65 (m, 6H), 3.61 – 3.47 (m, 8H), 3.41 (q, *J* = 5.8 Hz, 2H), 2.54 (s, 2H), 2.40 (s, 3H), 2.10 – 2.01 (m, 1H), 1.88 (ddd, *J* = 13.1, 8.9, 4.5 Hz, 1H), 0.90 (s, 9H). HRMS (ESI-MS) calcd for $C_{54}H_{62}FN_{10}O_9S^+$ m/z $(M+H)^+$ 1045.4400, found: 1045.4402.

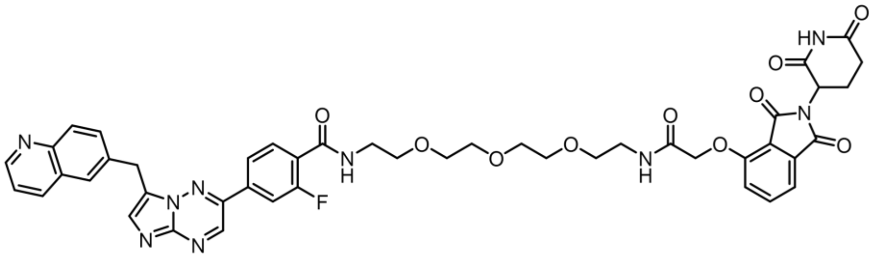

N-(1-((2-(2,6-dioxopiperidin-3-yl)-1,3-dioxoisoindolin-4-yl)oxy)-2-oxo-6,9,12-trioxa-3-azatetradecan-14-yl)-2-fluoro-4-(7-(quinolin-6-ylmethyl)imidazo[1,2-b][1,2,4]triazin-2-yl)benzamide (48-272)

Yield = 74%. $^1$H NMR (600 MHz, DMSO-$d_6$) δ 11.12 (s, 1H), 9.17 (s, 1H), 8.95 (dd, *J* = 4.6, 1.6 Hz, 1H), 8.63 – 8.54 (m, 1H), 8.43 (td, *J* = 5.7, 2.4 Hz, 1H), 8.07 (d, *J* = 1.8 Hz, 1H), 8.03 (d, *J* = 8.7 Hz, 1H), 8.00 (d, *J* = 6.7 Hz, 2H), 7.99 – 7.95 (m, 2H), 7.92 (dd, *J* = 8.8, 2.0 Hz, 1H), 7.76 (q, *J* = 8.1 Hz, 2H), 7.68 (dd, *J* = 8.4, 4.6 Hz, 1H), 7.44 (d, *J* = 7.3 Hz, 1H), 7.34 (d, *J* = 8.5 Hz, 1H), 5.07 (dd, *J* = 12.9, 5.5 Hz, 1H), 4.72 (s, 2H), 4.66 (s, 2H), 3.55 – 3.47 (m, 7H), 3.43 (dt, *J* = 14.0, 5.8 Hz, 4H), 3.29 (q, *J* = 5.7 Hz, 2H), 2.87 – 2.81 (m, 1H), 2.59 (dd, *J* = 17.3, 3.4 Hz, 1H), 2.54 (s, 2H), 2.02 (dtt, *J* = 13.0, 5.5, 2.3 Hz, 1H). HRMS (ESI-MS) calcd for $C_{45}H_{43}FN_9O_{10}^+$ m/z $(M+H)^+$ 888.3111, found: 888.3111.

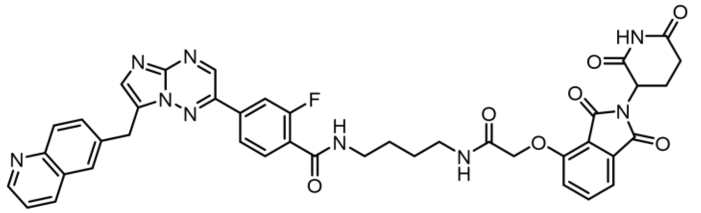

N-(4-(2-((2-(2,6-dioxopiperidin-3-yl)-1,3-dioxoisoindolin-4-yl)oxy)acetamido)butyl)-2-fluoro-4-(7-(quinolin-6-ylmethyl)imidazo[1,2-b][1,2,4]triazin-2-yl)benzamide (48-273)
Yield = 72%. $^1$H NMR (600 MHz, DMSO-$d_6$) δ 11.11 (s, 1H), 9.18 (s, 1H), 8.95 (dd, *J* = 4.6, 1.6 Hz, 1H), 8.61 – 8.56 (m, 1H), 8.48 (td, *J* = 5.7, 1.7 Hz, 1H), 8.10 – 7.90 (m, 8H), 7.79 (dd, *J* = 8.5, 7.3 Hz, 1H), 7.74 (t, *J* = 7.7 Hz, 1H), 7.69 (dd, *J* = 8.4, 4.6 Hz, 1H), 7.47 (d, *J* = 7.3 Hz, 1H), 7.38 (d, *J* = 8.5 Hz, 1H), 5.08 (dd, *J* = 12.9, 5.5 Hz, 1H), 4.76 (s, 2H), 4.67 (s, 2H), 3.26 (q, *J* = 6.3 Hz, 2H), 3.19 (q, *J* = 6.1, 5.7 Hz, 2H), 2.89 – 2.79 (m, 1H), 2.54 (s, 2H), 2.09 – 1.98 (m, 1H), 1.51 (dtt, *J* = 11.5, 8.3, 4.8 Hz, 4H). HRMS (ESI-MS) calcd for $C_{41}H_{35}FN_9O_7^+$ m/z $(M+H)^+$ 784.2638, found: 784.2636.

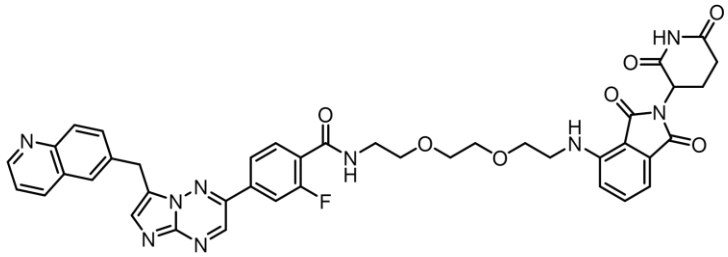

N-(2-(2-(2-((2-(2,6-dioxopiperidin-3-yl)-1,3-dioxoisoindolin-4-yl)amino)ethoxy)ethoxy)ethyl)-2-fluoro-4-(7-(quinolin-6-ylmethyl)imidazo[1,2-b][1,2,4]triazin-2-yl)benzamide (48-274)
Yield = 58%. $^1$H NMR (499 MHz, DMSO-$d_6$) δ 11.10 (s, 1H), 9.22 (s, 1H), 8.98 (dd, *J* = 4.5, 1.7 Hz, 1H), 8.55 (d, *J* = 8.5 Hz, 1H), 8.39 (dt, *J* = 5.7, 3.8 Hz, 1H), 8.10 – 7.98 (m, 5H), 7.92 (dd, *J* = 8.7, 2.0 Hz, 1H), 7.79 (t, *J* = 7.8 Hz, 1H), 7.67 (dd, *J* = 8.4, 4.5 Hz, 1H), 7.52 (dd, *J* = 8.5, 7.0 Hz, 1H), 7.10 (d, *J* = 8.6 Hz, 1H), 6.97 (d, *J* = 7.0 Hz, 1H), 6.59 (s, 1H), 5.05 (dd, *J* = 12.7, 5.4 Hz, 1H), 4.68 (s, 2H), 3.67 – 3.54 (m, 8H), 3.45 (q, *J* = 5.8 Hz, 4H), 2.93 – 2.82 (m, 1H), 2.63 – 2.52 (m, 2H), 2.02 (ddd, *J* = 12.0, 6.5, 4.0 Hz, 1H). HRMS (ESI-MS) calcd for $C_{41}H_{37}FN_9O_7^+$ m/z $(M+H)^+$ 786.2794, found: 786.2798.

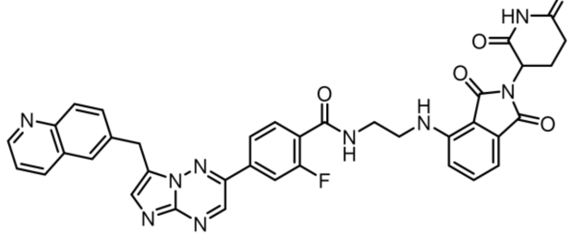

N-(2-((2-(2,6-dioxopiperidin-3-yl)-1,3-dioxoisoindolin-4-yl)amino)ethyl)-2-fluoro-4-(7-(quinolin-6-ylmethyl)imidazo[1,2-b][1,2,4]triazin-2-yl)benzamide (48-275)
Yield = 81%. $^1$H NMR (499 MHz, DMSO-$d_6$) δ 11.11 (s, 1H), 9.25 (s, 1H), 8.97 (dd, *J* = 4.5, 1.7 Hz, 1H), 8.69 (q, *J* = 5.0 Hz, 1H), 8.52 (d, *J* = 8.5 Hz, 1H), 8.14 – 7.98 (m, 5H), 7.91 (dd, *J* = 8.8, 2.0 Hz, 1H), 7.80 (t, *J* = 7.7 Hz, 1H), 7.71 – 7.56 (m, 2H), 7.27 (d, *J* = 8.7 Hz, 1H), 7.06 (d, *J* = 7.0 Hz, 1H), 6.89 – 6.79 (m, 1H), 5.07 (dd, *J* = 12.7, 5.5 Hz, 1H),

4.69 (s, 2H), 3.59 – 3.53 (m, 2H), 3.50 (q, *J* = 7.2, 6.2 Hz, 2H), 2.89 (ddd, *J* = 16.7, 13.4, 5.4 Hz, 1H), 2.66 – 2.56 (m, 2H), 2.03 (dp, *J* = 11.2, 3.6 Hz, 1H). HRMS (ESI-MS) calcd for $C_{37}H_{29}FN_9O_5^+$ m/z $(M+H)^+$ 698.2270, found: 698.2266.

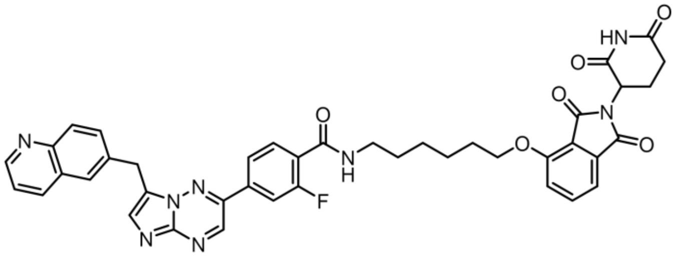

N-(6-((2-(2,6-dioxopiperidin-3-yl)-1,3-dioxoisoindolin-4-yl)oxy)hexyl)-2-fluoro-4-(7-(quinolin-6-ylmethyl)imidazo[1,2-b][1,2,4]triazin-2-yl)benzamide (48-276)
Yield = 69%. $^1$H NMR (499 MHz, DMSO-$d_6$) δ 11.11 (s, 1H), 9.24 (s, 1H), 8.96 (dd, *J* = 4.6, 1.6 Hz, 1H), 8.61 – 8.41 (m, 2H), 8.12 – 7.99 (m, 4H), 7.91 (dd, *J* = 8.8, 2.0 Hz, 1H), 7.84 – 7.74 (m, 2H), 7.66 (dd, *J* = 8.3, 4.5 Hz, 1H), 7.53 (d, *J* = 8.5 Hz, 1H), 7.44 (d, *J* = 7.3 Hz, 1H), 5.08 (dd, *J* = 12.8, 5.5 Hz, 1H), 4.68 (s, 2H), 4.23 (t, *J* = 6.4 Hz, 2H), 3.29 (q, *J* = 6.6 Hz, 2H), 2.94 – 2.84 (m, 1H), 2.58 (d, *J* = 30.4 Hz, 2H), 2.03 (dp, *J* = 11.1, 3.5 Hz, 1H), 1.79 (p, *J* = 6.7 Hz, 2H), 1.55 (dp, *J* = 22.7, 7.1 Hz, 4H), 1.43 (q, *J* = 7.8 Hz, 2H). HRMS (ESI-MS) calcd for $C_{41}H_{36}FN_8O_6^+$ m/z $(M+H)^+$ 755.2736, found: 755.2739.

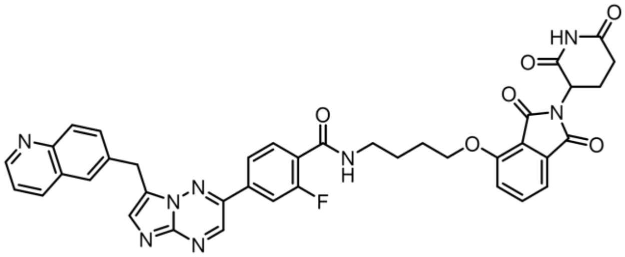

N-(4-((2-(2,6-dioxopiperidin-3-yl)-1,3-dioxoisoindolin-4-yl)oxy)butyl)-2-fluoro-4-(7-(quinolin-6-ylmethyl)imidazo[1,2-b][1,2,4]triazin-2-yl)benzamide (48-277)
Yield = 77%. $^1$H NMR (499 MHz, DMSO-$d_6$) δ 11.10 (s, 1H), 9.24 (s, 1H), 8.96 (dd, *J* = 4.5, 1.6 Hz, 1H), 8.53 (t, *J* = 5.6 Hz, 2H), 8.12 – 7.99 (m, 4H), 7.91 (dd, *J* = 8.7, 2.0 Hz, 1H), 7.86 – 7.71 (m, 2H), 7.65 (dd, *J* = 8.4, 4.5 Hz, 1H), 7.55 (d, *J* = 8.5 Hz, 1H), 7.46 (d, *J* = 7.2 Hz, 1H), 5.08 (dd, *J* = 12.7, 5.4 Hz, 1H), 4.68 (s, 2H), 4.28 (t, *J* = 6.3 Hz, 2H), 3.37 (q, *J* = 6.6 Hz, 2H), 2.92 – 2.83 (m, 1H), 2.60 (d, *J* = 3.4 Hz, 2H), 2.03 (ddd, *J* = 11.4, 6.3, 3.8 Hz, 1H), 1.86 (dq, *J* = 11.7, 6.4 Hz, 2H), 1.75 (q, *J* = 7.3 Hz, 2H). HRMS (ESI-MS) calcd for $C_{39}H_{32}FN_8O_6^+$ m/z $(M+H)^+$ 727.2423, found: 727.2421.

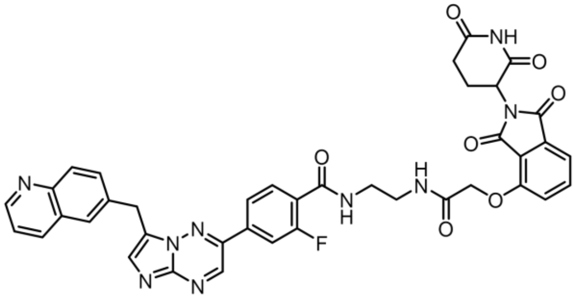

N-(2-(2-((2-(2,6-dioxopiperidin-3-yl)-1,3-dioxoisoindolin-4-yl)oxy)acetamido)ethyl)-2-fluoro-4-(7-(quinolin-6-ylmethyl)imidazo[1,2-b][1,2,4]triazin-2-yl)benzamide (48-278)
Yield = 78%. $^1$H NMR (499 MHz, DMSO-$d_6$) δ 11.12 (s, 1H), 9.24 (s, 1H), 8.94 (dd, *J* = 4.5, 1.7 Hz, 1H), 8.49 (s, 2H), 8.16 (t, *J* = 5.7 Hz, 1H), 8.10 – 7.94 (m, 5H), 7.89 (dd, *J* = 8.8, 2.0 Hz, 1H), 7.83 – 7.72 (m, 2H), 7.63 (dd, *J* = 8.4, 4.4 Hz, 1H), 7.46 (d, *J* = 7.2 Hz, 1H), 7.40 (d, *J* = 8.6 Hz, 1H), 5.10 (dd, *J* = 12.7, 5.4 Hz, 1H), 4.80 (s, 2H), 4.68 (s, 2H), 3.38 (dt, *J* = 10.6, 5.7 Hz, 3H), 2.94 – 2.82 (m, 2H), 2.31 – 2.58 (m, 2H), 2.01 (ddd, *J* = 10.9, 6.0, 3.3 Hz, 1H). HRMS (ESI-MS) calcd for $C_{39}H_{31}FN_9O_7^+$ m/z $(M+H)^+$ 756.2325, found: 756.2327.

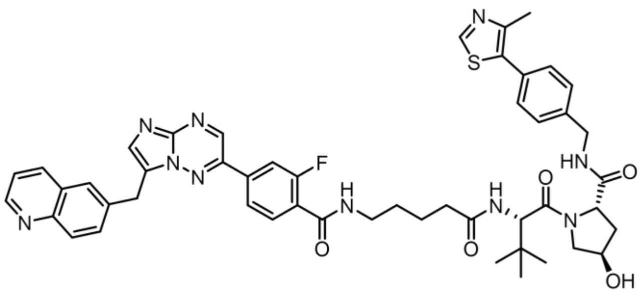

(2S,4R)-1-((S)-2-(5-(2-fluoro-4-(7-(quinolin-6-ylmethyl)imidazo[1,2-b][1,2,4]triazin-2-yl)benzamido)pentanamido)-3,3-dimethylbutanoyl)-4-hydroxy-N-(4-(4-methylthiazol-5-yl)benzyl)pyrrolidine-2-carboxamide (48-279)
Yield = 61%. $^1$H NMR (499 MHz, DMSO-$d_6$) δ 9.24 (s, 1H), 8.98 (d, *J* = 3.5 Hz, 2H), 8.57 (t, *J* = 6.2 Hz, 2H), 8.46 (t, *J* = 5.2 Hz, 1H), 8.11 – 8.01 (m, 5H), 7.95 – 7.86 (m, 2H), 7.77 (t, *J* = 7.7 Hz, 1H), 7.69 (dd, *J* = 8.4, 4.5 Hz, 1H), 7.46 – 7.34 (m, 4H), 4.68 (s, 2H), 4.56 (d, *J* = 9.4 Hz, 1H), 4.47 – 4.41 (m, 2H), 4.38 – 4.34 (m, 2H), 4.22 (dd, *J* = 15.9, 5.4 Hz, 2H), 3.27 (q, *J* = 6.2 Hz, 2H), 2.40 – 2.26 (m, 2H), 2.17 (dd, *J* = 14.5, 7.6 Hz, 2H), 2.09 – 1.98 (m, 2H), 1.91 (ddd, *J* = 14.4, 8.6, 4.6 Hz, 2H), 1.62 – 1.50 (m, 4H), 0.95 (s, 9H). HRMS (ESI-MS) calcd for $C_{49}H_{52}FN_{10}O_5S^+$ m/z $(M+H)^+$ 911.3821, found: 911.3816.

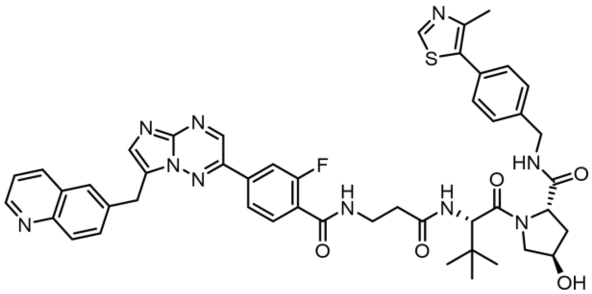

(2S,4R)-1-((S)-2-(3-(2-fluoro-4-(7-(quinolin-6-ylmethyl)imidazo[1,2-b][1,2,4]triazin-2-yl)benzamido)propanamido)-3,3-dimethylbutanoyl)-4-hydroxy-N-(4-(4-methylthiazol-5-yl)benzyl)pyrrolidine-2-carboxamide (48-280)
Yield = 74%. $^1$H NMR (499 MHz, DMSO-$d_6$) δ 9.24 (s, 1H), 9.02 – 8.95 (m, 2H), 8.58 (t, *J* = 6.3 Hz, 2H), 8.43 (q, *J* = 5.3 Hz, 1H), 8.12 – 8.01 (m, 6H), 7.93 (dd, *J* = 8.7, 2.0 Hz, 1H), 7.81 (t, *J* = 7.8 Hz, 1H), 7.69 (dd, *J* = 8.3, 4.5 Hz, 1H), 7.45 – 7.34 (m, 4H), 4.69 (s, 2H), 4.59 (s, 1H), 4.48 – 4.42 (m, 2H), 4.39 – 4.35 (m, 2H), 4.24 – 4.20 (m, 2H), 3.70 – 3.64 (m, 3H), 3.49 (p, *J* = 6.7 Hz, 3H), 2.59 (dt, *J* = 15.0, 7.5 Hz, 2H), 2.05 (t, *J* = 10.5 Hz, 1H), 1.92 (ddd, *J* = 11.0, 8.5, 4.6 Hz, 1H), 0.95 (s, 9H). HRMS (ESI-MS) calcd for $C_{47}H_{48}FN_{10}O_5S^+$ m/z $(M+H)^+$ 883.3508, found: 883.3503.

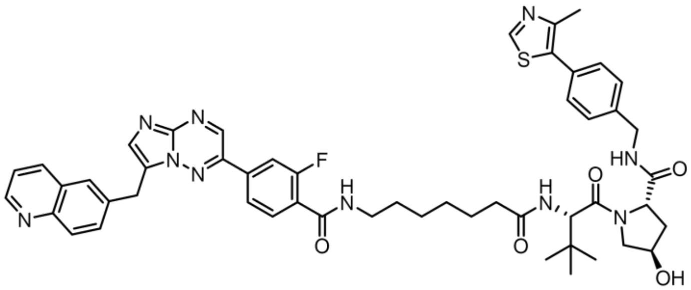

(2S,4R)-1-((S)-2-(7-(2-fluoro-4-(7-(quinolin-6-ylmethyl)imidazo[1,2-b][1,2,4]triazin-2-yl)benzamido)heptanamido)-3,3-dimethylbutanoyl)-4-hydroxy-N-(4-(4-methylthiazol-5-yl)benzyl)pyrrolidine-2-carboxamide (48-281)

Yield = 75%. $^{1}$H NMR (499 MHz, DMSO-$d_6$) δ 9.24 (s, 1H), 8.98 (d, *J* = 3.6 Hz, 2H), 8.56 (t, *J* = 6.1 Hz, 2H), 8.48 – 8.40 (m, 1H), 8.09 – 8.01 (m, 5H), 7.92 (dd, *J* = 8.8, 2.0 Hz, 1H), 7.86 (d, *J* = 9.4 Hz, 1H), 7.77 (t, *J* = 7.7 Hz, 1H), 7.68 (dd, *J* = 8.4, 4.5 Hz, 1H), 7.45 – 7.35 (m, 4H), 4.68 (s, 2H), 4.56 (d, *J* = 9.4 Hz, 1H), 4.43 (dd, *J* = 9.5, 6.6 Hz, 2H), 4.37 – 4.33 (m, 2H), 4.24 (d, *J* = 5.6 Hz, 2H), 3.67 (d, *J* = 8.4 Hz, 2H), 3.26 (q, *J* = 6.7 Hz, 2H), 2.28 (dt, *J* = 14.6, 7.5 Hz, 2H), 2.16 – 2.12 (m, 1H), 2.03 (d, *J* = 8.6 Hz, 1H), 1.93 – 1.88 (m, 1H), 1.52 (t, *J* = 7.9 Hz, 4H), 1.30 (d, *J* = 12.6 Hz, 5H), 0.95 (s, 9H). HRMS (ESI-MS) calcd for $C_{51}H_{56}FN_{10}O_5S^+$ m/z $(M+H)^+$ 939.4134, found: 939.4134.

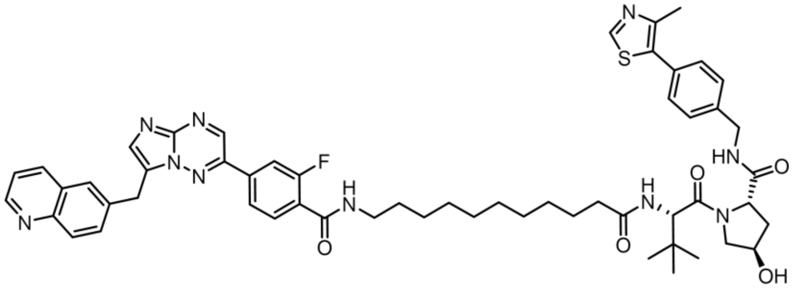

(2S,4R)-1-((S)-2-(11-(2-fluoro-4-(7-(quinolin-6-ylmethyl)imidazo[1,2-b][1,2,4]triazin-2-yl)benzamido)undecanamido)-3,3-dimethylbutanoyl)-4-hydroxy-N-(4-(4-methylthiazol-5-yl)benzyl)pyrrolidine-2-carboxamide (48-282)

Yield = 68%. $^{1}$H NMR (499 MHz, DMSO-$d_6$) δ 9.24 (s, 1H), 8.98 (d, *J* = 4.3 Hz, 2H), 8.56 (t, *J* = 5.9 Hz, 2H), 8.44 (t, *J* = 5.4 Hz, 1H), 8.11 – 8.00 (m, 5H), 7.92 (dd, *J* = 8.7, 2.0 Hz, 1H), 7.84 (d, *J* = 9.4 Hz, 1H), 7.76 (t, *J* = 7.7 Hz, 1H), 7.68 (dd, *J* = 8.4, 4.5 Hz, 1H), 7.47 – 7.35 (m, 4H), 4.68 (s, 2H), 4.55 (d, *J* = 9.4 Hz, 1H), 4.47 – 4.40 (m, 2H), 4.37 – 4.33 (m, 2H), 4.22 (dd, *J* = 15.9, 5.4 Hz, 2H), 3.71 – 3.64 (m, 5H), 3.26 (q, *J* = 6.7 Hz, 2H), 2.44 (s, 3H), 2.27 (dt, *J* = 14.7, 7.6 Hz, 1H), 2.12 (dd, *J* = 8.0, 6.1 Hz, 1H), 2.03 (t, *J* = 10.2 Hz, 1H), 1.90 (ddd, *J* = 12.8, 8.6, 4.6 Hz, 1H), 1.58 – 1.43 (m, 4H), 1.29 (d, *J* = 24.4 Hz, 8H), 0.93 (s, 9H). HRMS (ESI-MS) calcd for $C_{55}H_{64}FN_{10}O_5S^+$ m/z $(M+H)^+$ 995.4760, found: 995.4762.

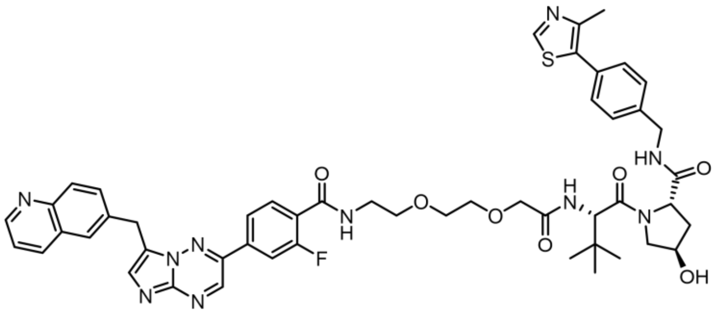

(2S,4R)-1-((S)-12-(tert-butyl)-1-(2-fluoro-4-(7-(quinolin-6-ylmethyl)imidazo[1,2-b][1,2,4]triazin-2-yl)phenyl)-1,10-dioxo-5,8-dioxa-2,11-diazatridecan-13-oyl)-4-hydroxy-N-(4-(4-methylthiazol-5-yl)benzyl)pyrrolidine-2-carboxamide (48-283)

Yield = 56%. $^1$H NMR (499 MHz, DMSO-$d_6$) δ 9.24 (s, 1H), 8.99 – 8.93 (m, 2H), 8.57 (t, *J* = 6.0 Hz, 1H), 8.53 (d, *J* = 8.3 Hz, 1H), 8.47 (q, *J* = 5.2 Hz, 1H), 8.12 – 7.99 (m, 5H), 7.91 (dd, *J* = 8.7, 2.0 Hz, 1H), 7.79 (t, *J* = 7.8 Hz, 1H), 7.66 (dd, *J* = 8.3, 4.5 Hz, 1H), 7.45 (d, *J* = 9.6 Hz, 1H), 7.39 (s, 4H), 4.68 (s, 2H), 4.58 (d, *J* = 9.6 Hz, 1H), 4.48 – 4.33 (m, 4H), 4.26 (dd, *J* = 15.7, 5.6 Hz, 2H), 4.00 (s, 2H), 3.66 – 3.59 (m, 7H), 3.47 (q, *J* = 5.9 Hz, 2H), 2.43 (s, 3H), 2.09 – 2.01 (m, 1H), 1.90 (ddd, *J* = 13.0, 8.5, 4.5 Hz, 1H), 0.94 (s, 9H). HRMS (ESI-MS) calcd for $C_{50}H_{54}FN_{10}O_7S^+$ m/z (M+H)$^+$ 957.3876, found: 957.3875.

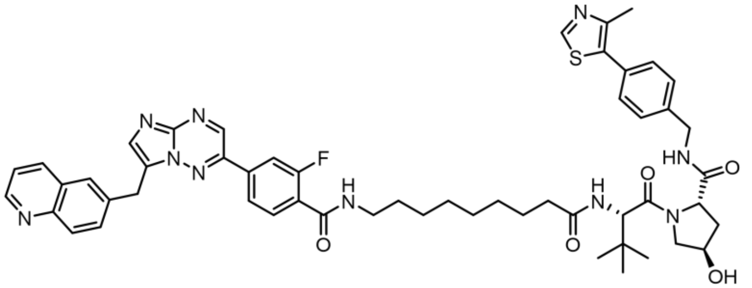

(2S,4R)-1-((S)-2-(9-(2-fluoro-4-(7-(quinolin-6-ylmethyl)imidazo[1,2-b][1,2,4]triazin-2-yl)benzamido)nonanamido)-3,3-dimethylbutanoyl)-4-hydroxy-N-(4-(4-methylthiazol-5-yl)benzyl)pyrrolidine-2-carboxamide (48-284)

Yield = 53%. $^1$H NMR (600 MHz, DMSO-$d_6$) δ 9.26 (s, 1H), 9.06 (dd, *J* = 4.7, 1.6 Hz, 1H), 8.99 (s, 1H), 8.70 (d, *J* = 8.3 Hz, 1H), 8.57 (t, *J* = 6.1 Hz, 1H), 8.44 (td, *J* = 5.6, 1.6 Hz, 1H), 8.14 (d, *J* = 1.9 Hz, 1H), 8.10 (d, *J* = 8.7 Hz, 1H), 8.06 – 8.02 (m, 3H), 7.99 (dd, *J* = 8.8, 1.9 Hz, 1H), 7.86 (d, *J* = 9.3 Hz, 1H), 7.77 (dt, *J* = 9.0, 6.2 Hz, 2H), 7.44 – 7.36 (m, 4H), 4.70 (s,2H), 4.55 (d, *J* = 9.4 Hz, 1H), 4.46 – 4.40 (m, 2H), 4.35 (tt, *J* = 4.6, 2.6 Hz, 1H), 4.22 (dd, *J* = 15.8, 5.5 Hz, 1H), 3.66 (qd, *J* = 9.8, 9.0, 2.9 Hz, 2H), 3.26 (q, *J* = 6.6 Hz, 2H), 2.44 (s, 3H), 2.31 – 2.24 (m, 1H), 2.12 (ddd, *J* = 14.2, 8.2, 6.2 Hz, 1H), 2.06 – 2.01 (m, 1H), 1.91 (ddd, *J* = 12.9, 8.6, 4.6 Hz, 1H), 1.56 – 1.43 (m, 4H), 1.37 – 1.21 (m, 9H), 0.94 (s, 9H). $^{13}$C NMR (151 MHz, DMSO) δ 172.05, 171.91, 169.67, 162.88, 160.06, 158.56, 158.41, 158.31, 158.07, 157.83, 151.44, 147.62, 144.04, 141.23, 140.98, 139.48, 137.29, 136.13, 134.45, 133.19, 131.14, 130.86, 129.56, 128.59, 127.60, 127.38, 125.93, 122.81, 121.93, 116.49, 114.64, 114.56, 114.47, 68.83, 58.66, 56.32, 56.23, 41.61, 39.90, 39.77, 39.63, 39.49, 39.35, 39.21, 39.07, 37.94, 35.19, 34.84, 28.87, 28.72, 28.64, 28.62, 26.35, 25.42, 15.90. HRMS (ESI-MS) calcd for $C_{53}H_{60}FN_{10}O_5S^+$ m/z (M+H)$^+$ 967.4447, found: 967.4449.

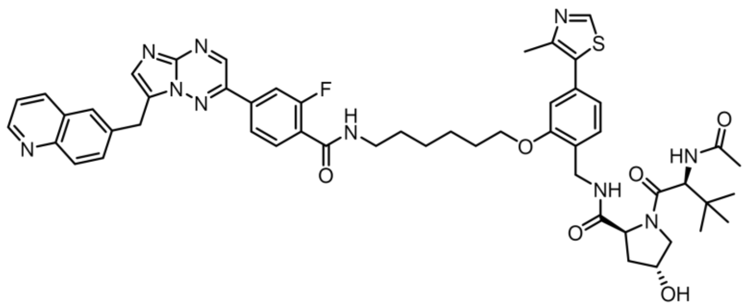

(2S,4R)-1-((S)-2-acetamido-3,3-dimethylbutanoyl)-N-(2-((6-(2-fluoro-4-(7-(quinolin-6-ylmethyl)imidazo[1,2-b][1,2,4]triazin-2-yl)benzamido)hexyl)oxy)-4-(4-methylthiazol-5-yl)benzyl)-4-hydroxypyrrolidine-2-carboxamide (48-285)

Yield = 74%. $^{1}$H NMR (499 MHz, DMSO-$d_6$) δ 9.24 (s, 1H), 8.98 (d, *J* = 7.3 Hz, 2H), 8.56 (d, *J* = 8.3 Hz, 1H), 8.46 (t, *J* = 5.9 Hz, 2H), 8.12 – 7.99 (m, 5H), 7.99 – 7.88 (m, 2H), 7.77 (t, *J* = 7.9 Hz, 1H), 7.68 (dd, *J* = 8.4, 4.5 Hz, 1H), 7.47 (d, *J* = 7.8 Hz, 1H), 7.00 (d, *J* = 1.7 Hz, 1H), 6.90 (dd, *J* = 7.8, 1.6 Hz, 1H), 4.69 (s, 2H), 4.54 (d, *J* = 9.4 Hz, 1H), 4.47 (t, *J* = 8.0 Hz, 1H), 4.37 – 4.32 (m, 2H), 4.16 (dd, *J* = 16.6, 5.5 Hz, 2H), 4.06 (d, *J* = 6.4 Hz, 2H), 3.67 – 3.63 (m, 3H), 3.29 (q, *J* = 6.6 Hz, 2H), 2.46 (s, 3H), 2.06 – 2.02 (m, 1H), 1.89 (s, 3H), 1.78 (q, *J* = 6.7, 6.2 Hz, 2H), 1.63 – 1.54 (m , 2H) 1.58 (p, *J* = 7.1 Hz, 2H), 1.51 (q, *J* = 7.5, 7.0 Hz, 2H), 1.43 (q, *J* = 7.9 Hz, 2H), 0.92 (s, 9H). HRMS (ESI-MS) calcd for $C_{52}H_{58}FN_{10}O_6S^+$ m/z (M+H)$^+$ 969.4240, found: 969.4240.

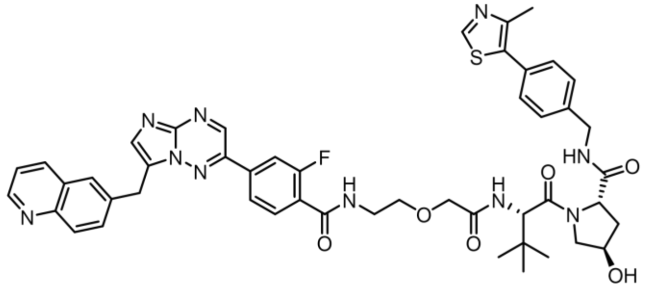

(2S,4R)-1-((S)-2-(2-(2-(2-fluoro-4-(7-(quinolin-6-ylmethyl)imidazo[1,2-b][1,2,4]triazin-2-yl)benzamido)ethoxy)acetamido)-3,3-dimethylbutanoyl)-4-hydroxy-N-(4-(4-methylthiazol-5-yl)benzyl)pyrrolidine-2-carboxamide (48-286)

Yield = 63%. $^{1}$H NMR (499 MHz, DMSO-$d_6$) δ 9.22 (s, 1H), 9.01 – 8.96 (m, 1H), 8.94 (s, 1H), 8.58 (dt, *J* = 13.5, 7.0 Hz, 3H), 8.10 – 7.99 (m, 5H), 7.92 (dd, *J* = 8.7, 2.1 Hz, 1H), 7.84 (t, *J* = 7.8 Hz, 1H), 7.69 (dd, *J* = 8.3, 4.5 Hz, 1H), 7.49 (d, *J* = 9.5 Hz, 1H), 7.37 (q, *J* = 8.4 Hz, 4H), 4.66 (s, 2H), 4.58 (d, *J* = 9.5 Hz, 1H), 4.50 – 4.33 (m, 3H), 4.25 (dd, *J* = 15.8, 5.6 Hz, 3H), 4.03 (d, *J* = 3.7 Hz, 3H), 3.51 – 3.44 (m, 2H), 2.40 (s, 3H), 2.11 – 2.00 (m, 2H), 1.91 (ddd, *J* = 13.1, 8.8, 4.6 Hz, 2H), 0.94 (s, 9H). HRMS (ESI-MS) calcd for $C_{48}H_{50}FN_{10}O_6S^+$ m/z (M+H)$^+$ 913.3614, found: 913.3610.

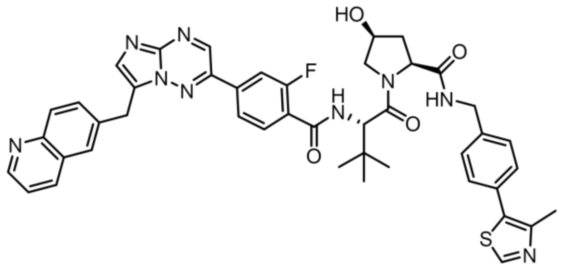

(2S,4S)-1-((S)-2-(2-fluoro-4-(7-(quinolin-6-ylmethyl)imidazo[1,2-b][1,2,4]triazin-2-yl)benzamido)-3,3-dimethylbutanoyl)-4-hydroxy-N-(4-(4-methylthiazol-5-yl)benzyl)pyrrolidine-2-carboxamide (48-287)

Yield = 68%. $^1$H NMR (499 MHz, DMSO-$d_6$) δ 9.25 (s, 1H), 9.05 – 8.91 (m, 2H), 8.69 (t, $J$ = 6.1 Hz, 1H), 8.56 (d, $J$ = 8.5 Hz, 1H), 8.36 (dd, $J$ = 8.6, 3.6 Hz, 1H), 8.16 – 7.97 (m, 5H), 7.92 (dd, $J$ = 8.7, 2.0 Hz, 1H), 7.78 (t, $J$ = 7.8 Hz, 1H), 7.68 (dd, $J$ = 8.3, 4.5 Hz, 1H), 7.51 – 7.32 (m, 4H), 4.69 (d, $J$ = 5.2 Hz, 2H), 4.49 – 4.40 (m, 2H), 4.34 – 4.29 (m, 2H), 4.02 (d, $J$ = 5.7 Hz, 2H), 3.53 (dd, $J$ = 10.1, 5.4 Hz, 2H), 2.45 (s, 3H), 2.39 (dd, $J$ = 6.4, 2.3 Hz, 1H), 1.78 (dt, $J$ = 12.3, 6.1 Hz, 1H), 1.06 (s, 9H). HRMS (ESI-MS) calcd for $C_{44}H_{43}FN_9O_4S^+$ m/z $(M+H)^+$ 812.3137, found: 812.3141.

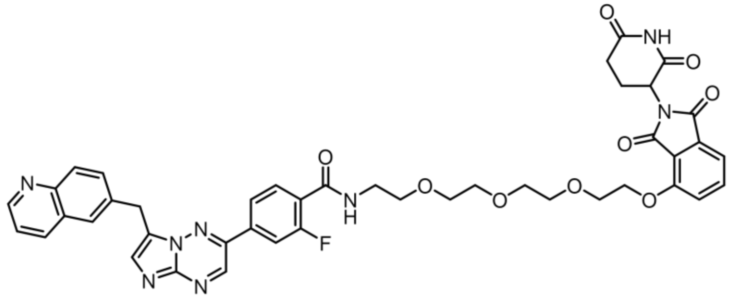

N-(2-(2-(2-(2-((2-(2,6-dioxopiperidin-3-yl)-1,3-dioxoisoindolin-4-yl)oxy)ethoxy)ethoxy)ethoxy)ethyl)-2-fluoro-4-(7-(quinolin-6-ylmethyl)imidazo[1,2-b][1,2,4]triazin-2-yl)benzamide (48-288)

Yield = 70%. $^1$H NMR (499 MHz, DMSO-$d_6$) δ 11.11 (s, 1H), 9.24 (s, 1H), 8.99 (dd, $J$ = 4.5, 1.7 Hz, 1H), 8.57 (d, $J$ = 8.3 Hz, 1H), 8.44 (td, $J$ = 5.8, 2.4 Hz, 1H), 8.10 – 8.00 (m, 5H), 7.92 (dd, $J$ = 8.7, 2.0 Hz, 1H), 7.83 – 7.73 (m, 2H), 7.69 (dd, $J$ = 8.3, 4.6 Hz, 1H), 7.50 (d, $J$ = 8.5 Hz, 1H), 7.42 (d, $J$ = 7.2 Hz, 1H), 5.08 (dd, $J$ = 12.8, 5.4 Hz, 1H), 4.68 (s, 2H), 4.32 (dd, $J$ = 5.6, 3.6 Hz, 2H), 3.82 – 3.77 (m, 2H), 3.65 (dd, $J$ = 5.9, 3.7 Hz, 2H), 3.56 (d, $J$ = 6.3 Hz, 8H), 3.44 (q, $J$ = 5.8 Hz, 2H), 2.94 – 2.83 (m, 1H), 2.59 (d, $J$ = 19.7 Hz, 2H), 2.03 (ddd, $J$ = 11.4, 5.9, 3.5 Hz, 1H). HRMS (ESI-MS) calcd for $C_{43}H_{40}FN_8O_9^+$ m/z $(M+H)^+$ 831.2897, found: 831.2899.

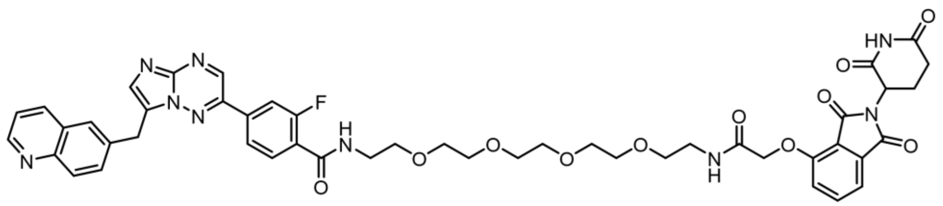

N-(1-((2-(2,6-dioxopiperidin-3-yl)-1,3-dioxoisoindolin-4-yl)oxy)-2-oxo-6,9,12,15-tetraoxa-3-azaheptadecan-17-yl)-2-fluoro-4-(7-(quinolin-6-ylmethyl)imidazo[1,2-b][1,2,4]triazin-2-yl)benzamide (48-289)

Yield = 59%. $^1$H NMR (499 MHz, DMSO-$d_6$) δ 11.12 (s, 1H), 9.24 (s, 1H), 8.98 (dd, $J$ = 4.5, 1.7 Hz, 1H), 8.55 (d, $J$ = 8.2 Hz, 1H), 8.45 (dt, $J$ = 5.6, 3.7 Hz, 1H), 8.10 – 7.96 (m, 6H), 7.92 (dd, $J$ = 8.8, 2.0 Hz, 1H), 7.80 (dd, $J$ = 8.4, 7.2 Hz, 2H), 7.68 (dd, $J$ = 8.3, 4.5 Hz, 1H), 7.48 (d, $J$ = 7.2 Hz, 1H), 7.39 (d, $J$ = 8.6 Hz, 1H), 5.11 (dd, $J$ = 12.8, 5.4 Hz, 1H), 4.78 (s, 2H), 4.68 (s, 2H), 3.59 – 3.49 (m, 14H), 3.49 – 3.40 (m, 4H), 3.31 (q, $J$ = 5.7 Hz, 2H), 2.90 (ddd, $J$ = 17.4, 13.9, 5.5 Hz, 1H), 2.67 – 2.56 (m, 2H), 2.06 – 2.01 (m, 1H). HRMS (ESI-MS) calcd for $C_{47}H_{47}FN_9O_{11}^+$ m/z (M+H)$^+$ 932.3374, found: 932.3373.

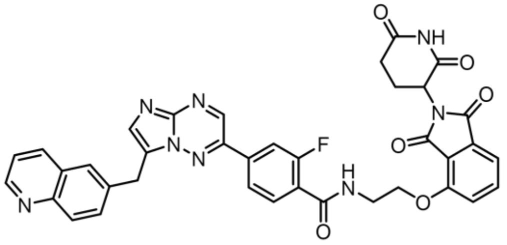

N-(2-((2-(2,6-dioxopiperidin-3-yl)-1,3-dioxoisoindolin-4-yl)oxy)ethyl)-2-fluoro-4-(7-(quinolin-6-ylmethyl)imidazo[1,2-b][1,2,4]triazin-2-yl)benzamide (48-290)

Yield = 55%. $^1$H NMR (499 MHz, DMSO-$d_6$) δ 11.13 (s, 1H), 9.25 (s, 1H), 8.99 (dd, $J$ = 4.6, 1.7 Hz, 1H), 8.65 (td, $J$ = 5.5, 2.4 Hz, 1H), 8.56 (d, $J$ = 8.4 Hz, 1H), 8.11 – 8.01 (m, 5H), 7.93 (dd, $J$ = 8.7, 2.0 Hz, 1H), 7.89 – 7.80 (m, 2H), 7.68 (dd, $J$ = 8.4, 4.5 Hz, 1H), 7.62 (d, $J$ = 8.6 Hz, 1H), 7.50 (d, $J$ = 7.2 Hz, 1H), 5.10 (dd, $J$ = 12.7, 5.4 Hz, 1H), 4.69 (s, 2H), 4.42 (t, $J$ = 6.0 Hz, 2H), 3.71 (q, $J$ = 5.8 Hz, 2H), 2.89 (ddd, $J$ = 17.0, 13.9, 5.4 Hz, 1H), 2.55 (s, 2H), 2.04 (ddq, $J$ = 10.5, 5.5, 3.2, 2.6 Hz, 1H). HRMS (ESI-MS) calcd for $C_{37}H_{28}FN_8O_6^+$ m/z (M+H)$^+$ 699.2110, found: 699.2110.

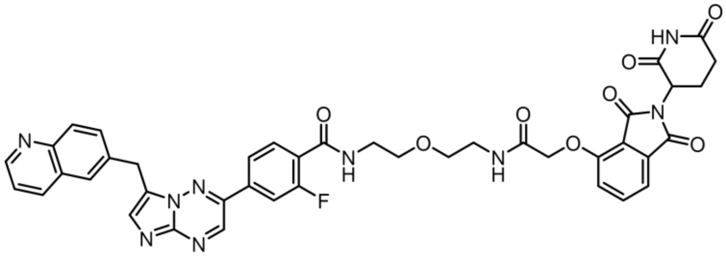

N-(2-(2-(2-((2-(2,6-dioxopiperidin-3-yl)-1,3-dioxoisoindolin-4-yl)oxy)acetamido)ethoxy)ethyl)-2-fluoro-4-(7-(quinolin-6-ylmethyl)imidazo[1,2-b][1,2,4]triazin-2-yl)benzamide (48-292)

Yield = 66%. $^1$H NMR (499 MHz, DMSO-$d_6$) δ 11.12 (s, 1H), 9.21 (s, 1H), 8.97 (dd, $J$ = 4.6, 1.7 Hz, 1H), 8.54 (d, $J$ = 8.4 Hz, 1H), 8.38 (dt, $J$ = 5.4, 3.6 Hz, 1H), 8.09 – 8.02 (m, 2H), 8.05 – 7.96 (m, 4H), 7.91 (dd, $J$ = 8.7, 2.0 Hz, 1H), 7.82 – 7.71 (m, 2H), 7.67 (dd, $J$ = 8.4, 4.5 Hz, 1H), 7.39 (dd, $J$ = 23.2, 7.9 Hz, 2H), 5.12 (dd, $J$ = 12.8, 5.4 Hz, 1H), 4.78 (s, 2H), 4.68 (s, 2H), 3.55 (dt, $J$ = 24.0, 5.8 Hz, 4H), 3.47 (q, $J$ = 5.8 Hz, 2H), 3.36 (q, $J$ = 5.8 Hz, 2H), 2.90 (ddd, $J$ = 16.7, 13.7, 5.4 Hz, 1H), 2.56 – 2.56 (m, 2H), 2.08 – 2.01 (m, 1H). HRMS (ESI-MS) calcd for $C_{41}H_{35}FN_9O_8^+$ m/z (M+H)$^+$ 800.2587, found: 800.2587.

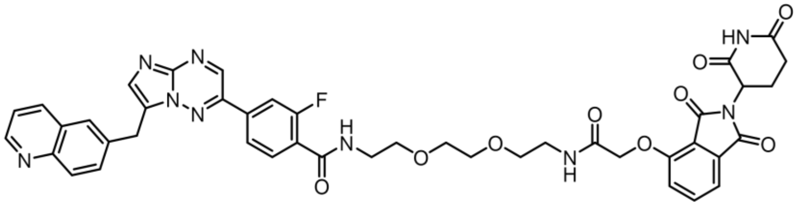

N-(2-(2-(2-(2-((2-(2,6-dioxopiperidin-3-yl)-1,3-dioxoisoindolin-4-yl)oxy)acetamido)ethoxy)ethoxy)ethyl)-2-fluoro-4-(7-(quinolin-6-ylmethyl)imidazo[1,2-b][1,2,4]triazin-2-yl)benzamide (48-293)

Yield = 90%. $^1$H NMR (499 MHz, DMSO-$d_6$) δ 11.12 (s, 1H), 9.24 (s, 1H), 8.97 (dd, *J* = 4.5, 1.7 Hz, 1H), 8.54 (d, *J* = 8.4 Hz, 1H), 8.45 (dt, *J* = 7.5, 3.7 Hz, 1H), 8.09 – 7.98 (m, 6H), 7.91 (dd, *J* = 8.7, 2.0 Hz, 1H), 7.83 – 7.75 (m, 2H), 7.67 (dd, *J* = 8.4, 4.5 Hz, 1H), 7.47 (d, *J* = 7.2 Hz, 1H), 7.38 (d, *J* = 8.6 Hz, 1H), 5.11 (dd, *J* = 12.8, 5.4 Hz, 1H), 4.78 (s, 2H), 4.68 (s, 2H), 3.57 (qd, *J* = 5.5, 2.5 Hz, 6H), 3.47 (dt, *J* = 21.8, 5.8 Hz, 4H), 3.33 (q, *J* = 5.7 Hz, 2H), 2.90 (ddd, *J* = 17.3, 13.8, 5.4 Hz, 1H), 2.64 – 2.53 (m, 2H), 2.04 (dp, *J* = 10.6, 3.4 Hz, 1H). HRMS (ESI-MS) calcd for $C_{43}H_{39}FN_9O_9^+$ m/z $(M+H)^+$ 844.2849, found: 844.2847.

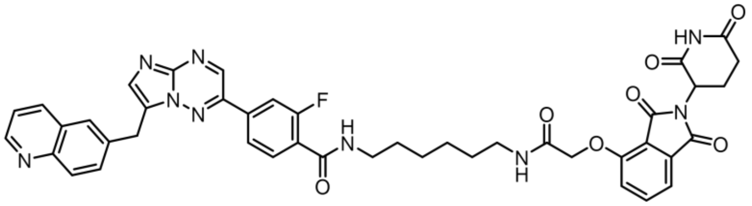

N-(6-(2-((2-(2,6-dioxopiperidin-3-yl)-1,3-dioxoisoindolin-4-yl)oxy)acetamido)hexyl)-2-fluoro-4-(7-(quinolin-6-ylmethyl)imidazo[1,2-b][1,2,4]triazin-2-yl)benzamide (48-294)

Yield = 87%. $^1$H NMR (499 MHz, DMSO-$d_6$) δ 11.12 (s, 1H), 9.24 (s, 1H), 8.95 (dd, *J* = 4.5, 1.7 Hz, 1H), 8.51 (d, *J* = 8.4 Hz, 1H), 8.47 – 8.41 (m, 1H), 8.04 (dd, *J* = 12.3, 3.7 Hz, 5H), 7.96 (t, *J* = 5.7 Hz, 1H), 7.90 (dd, *J* = 8.7, 2.0 Hz, 1H), 7.85 – 7.73 (m, 2H), 7.65 (dd, *J* = 8.3, 4.4 Hz, 1H), 7.49 (d, *J* = 7.2 Hz, 1H), 7.40 (d, *J* = 8.5 Hz, 1H), 5.13 (dd, *J* = 12.8, 5.4 Hz, 1H), 4.78 (s, 2H), 4.68 (s, 2H), 3.26 (q, *J* = 6.6 Hz, 2H), 3.21 – 3.13 (m, 2H), 2.90 (ddd, *J* = 16.9, 13.8, 5.4 Hz, 1H), 2.64 – 2.52 (m, 2H), 2.07 – 2.00 (m, 1H), 1.49 (dq, *J* = 27.1, 6.4, 6.0 Hz, 4H), 1.37 – 1.30 (m, 4H). HRMS (ESI-MS) calcd for $C_{43}H_{39}FN_9O_7^+$ m/z $(M+H)^+$ 812.2951, found: 812.2953.

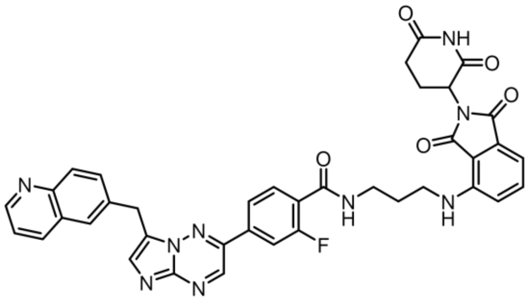

N-(3-((2-(2,6-dioxopiperidin-3-yl)-1,3-dioxoisoindolin-4-yl)amino)propyl)-2-fluoro-4-(7-(quinolin-6-ylmethyl)imidazo[1,2-b][1,2,4]triazin-2-yl)benzamide (48-295)

Yield = 65%. $^1$H NMR (499 MHz, DMSO-$d_6$) δ 11.10 (s, 1H), 9.25 (s, 1H), 8.98 (dd, *J* = 4.6, 1.7 Hz, 1H), 8.61 – 8.53 (m, 2H), 8.10 – 8.01 (m, 5H), 7.92 (dd, *J* = 8.8, 2.0 Hz, 1H),

7.81 (t, $J$ = 7.6 Hz, 1H), 7.68 (dd, $J$ = 8.3, 4.5 Hz, 1H), 7.60 (dd, $J$ = 8.6, 7.1 Hz, 1H), 7.14 (d, $J$ = 8.7 Hz, 1H), 7.04 (d, $J$ = 7.0 Hz, 1H), 6.80 – 6.75 (m, 1H), 5.06 (dd, $J$ = 12.7, 5.4 Hz, 1H), 4.69 (s, 2H), 3.39 (dt, $J$ = 19.3, 6.3 Hz, 4H), 2.88 (ddd, $J$ = 16.6, 13.6, 5.3 Hz, 1H), 2.63 – 2.54 (m, 2H), 2.03 (ddt, $J$ = 12.9, 5.5, 3.0 Hz, 1H), 1.83 (p, $J$ = 6.7 Hz, 2H). HRMS (ESI-MS) calcd for $C_{38}H_{31}FN_9O_5^+$ m/z $(M+H)^+$ 712.2427, found: 712.2425.

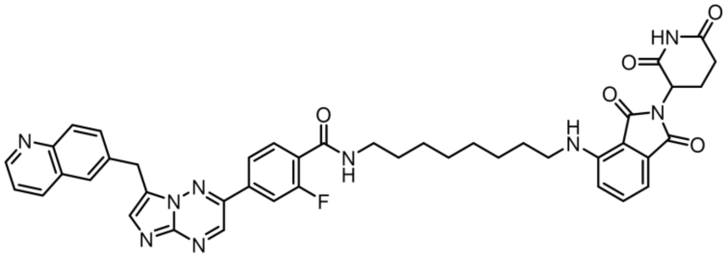

N-(8-((2-(2,6-dioxopiperidin-3-yl)-1,3-dioxoisoindolin-4-yl)amino)octyl)-2-fluoro-4-(7-(quinolin-6-ylmethyl)imidazo[1,2-b][1,2,4]triazin-2-yl)benzamide (48-296)
Yield = 60%. $^1$H NMR (499 MHz, DMSO-$d_6$) δ 11.09 (s, 1H), 9.24 (s, 1H), 8.98 (dd, $J$ = 4.6, 1.6 Hz, 1H), 8.56 (d, $J$ = 7.9 Hz, 1H), 8.47 – 8.40 (m, 1H), 8.05 (dd, $J$ = 16.8, 8.9 Hz, 5H), 7.92 (dd, $J$ = 8.7, 2.0 Hz, 1H), 7.76 (t, $J$ = 7.7 Hz, 1H), 7.68 (dd, $J$ = 8.4, 4.5 Hz, 1H), 7.57 (dd, $J$ = 8.6, 7.0 Hz, 1H), 7.09 (d, $J$ = 8.6 Hz, 1H), 7.01 (d, $J$ = 7.0 Hz, 1H), 6.55 – 6.51 (m, 1H), 5.05 (dd, $J$ = 12.8, 5.5 Hz, 1H), 4.69 (s, 2H), 3.27 (dt, $J$ = 13.1, 7.0 Hz, 4H), 2.94 – 2.83 (m, 1H), 2.67 – 2.50 (m, 2H), 2.03 (tt, $J$ = 7.4, 4.1 Hz, 1H), 1.56 (dt, $J$ = 28.3, 7.2 Hz, 4H), 1.34 (s, 7H). HRMS (ESI-MS) calcd for $C_{43}H_{41}FN_9O_5^+$ m/z $(M+H)^+$ 782.3209, found: 782.3211.

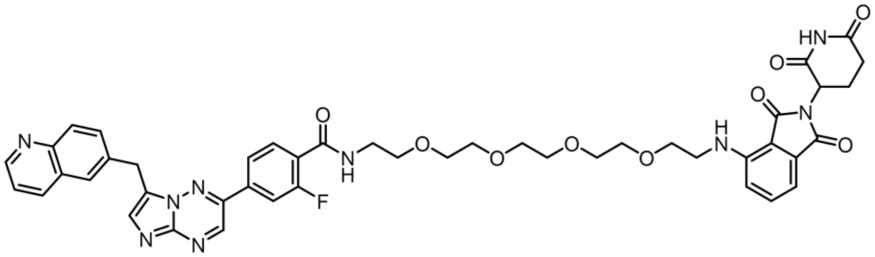

N-(14-((2-(2,6-dioxopiperidin-3-yl)-1,3-dioxoisoindolin-4-yl)amino)-3,6,9,12-tetraoxatetradecyl)-2-fluoro-4-(7-(quinolin-6-ylmethyl)imidazo[1,2-b][1,2,4]triazin-2-yl)benzamide (48-297)
Yield = 76%. $^1$H NMR (499 MHz, DMSO-$d_6$) δ 11.10 (s, 1H), 9.24 (s, 1H), 8.99 (dd, $J$ = 4.5, 1.7 Hz, 1H), 8.57 (d, $J$ = 8.4 Hz, 1H), 8.44 (td, $J$ = 5.6, 2.3 Hz, 1H), 8.10 – 8.00 (m, 5H), 7.92 (dd, $J$ = 8.7, 2.0 Hz, 1H), 7.80 (t, $J$ = 7.9 Hz, 1H), 7.69 (dd, $J$ = 8.3, 4.5 Hz, 1H), 7.55 (dd, $J$ = 8.6, 7.0 Hz, 1H), 7.11 (d, $J$ = 8.6 Hz, 1H), 7.01 (d, $J$ = 7.1 Hz, 1H), 6.60 – 6.55 (m, 1H), 5.05 (dd, $J$ = 12.7, 5.4 Hz, 1H), 4.68 (s, 2H), 3.60 (t, $J$ = 5.5 Hz, 2H), 3.58 – 3.47 (m, 14H), 3.44 (q, $J$ = 5.8 Hz, 4H), 2.89 (ddd, $J$ = 13.9, 11.7, 7.0 Hz, 1H), 2.63 – 2.52 (m, 2H), 2.07 – 1.98 (m, 1H). HRMS (ESI-MS) calcd for $C_{45}H_{45}FN_9O_9^+$ m/z $(M+H)^+$ 874.3314, found: 874.3319.

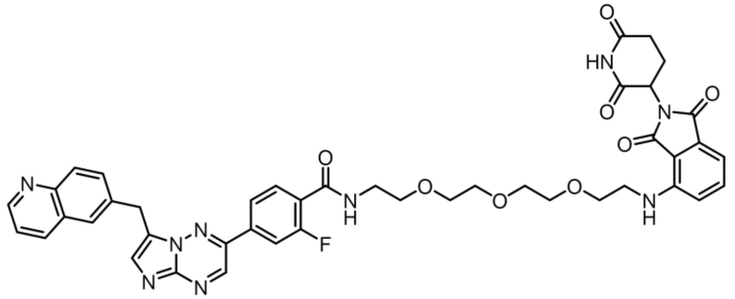

N-(2-(2-(2-(2-((2-(2,6-dioxopiperidin-3-yl)-1,3-dioxoisoindolin-4-yl)amino)ethoxy)ethoxy)ethoxy)ethyl)-2-fluoro-4-(7-(quinolin-6-ylmethyl)imidazo[1,2-b][1,2,4]triazin-2-yl)benzamide (48-298)

Yield = 61%. $^1$H NMR (499 MHz, DMSO-$d_6$) δ 11.10 (s, 1H), 9.24 (s, 1H), 8.99 (dd, $J$ = 4.6, 1.6 Hz, 1H), 8.57 (d, $J$ = 8.3 Hz, 1H), 8.46 – 8.39 (m, 1H), 8.10 – 8.00 (m, 5H), 7.93 (dd, $J$ = 8.8, 2.0 Hz, 1H), 7.80 (t, $J$ = 7.8 Hz, 1H), 7.69 (dd, $J$ = 8.4, 4.6 Hz, 1H), 7.54 (dd, $J$ = 8.6, 7.1 Hz, 1H), 7.10 (d, $J$ = 8.6 Hz, 1H), 7.00 (d, $J$ = 7.0 Hz, 1H), 6.60 – 6.55 (m, 1H), 5.05 (dd, $J$ = 12.7, 5.4 Hz, 1H), 4.68 (s, 2H), 3.61 (t, $J$ = 5.5 Hz, 2H), 3.56 (d, $J$ = 5.1 Hz, 10H), 3.44 (q, $J$ = 5.8 Hz, 4H), 2.89 (ddd, $J$ = 14.0, 11.5, 7.1 Hz, 1H), 2.63 – 2.54 (m, 2H), 2.07 – 1.98 (m, 1H). HRMS (ESI-MS) calcd for $C_{43}H_{41}FN_9O_8^+$ m/z $(M+H)^+$ 830.3057, found: 830.3056.

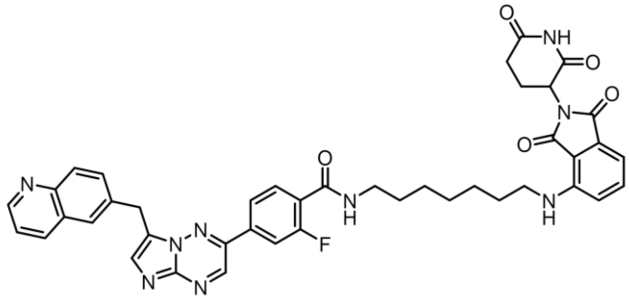

N-(7-((2-(2,6-dioxopiperidin-3-yl)-1,3-dioxoisoindolin-4-yl)amino)heptyl)-2-fluoro-4-(7-(quinolin-6-ylmethyl)imidazo[1,2-b][1,2,4]triazin-2-yl)benzamide (48-299)

Yield = 89%. $^1$H NMR (499 MHz, DMSO-$d_6$) δ 11.09 (s, 1H), 9.24 (s, 1H), 8.96 (dd, $J$ = 4.5, 1.6 Hz, 1H), 8.52 (d, $J$ = 8.4 Hz, 1H), 8.44 (t, $J$ = 5.6 Hz, 1H), 8.08 – 8.00 (m, 5H), 7.90 (dd, $J$ = 8.8, 2.0 Hz, 1H), 7.76 (t, $J$ = 7.8 Hz, 1H), 7.65 (dd, $J$ = 8.4, 4.5 Hz, 1H), 7.58 (dd, $J$ = 8.6, 7.0 Hz, 1H), 7.10 (d, $J$ = 8.6 Hz, 1H), 7.01 (d, $J$ = 7.1 Hz, 1H), 6.60 – 6.54 (m, 1H), 5.05 (dd, $J$ = 12.7, 5.4 Hz, 1H), 4.68 (s, 2H), 3.33 – 3.23 (m, 4H), 2.88 (ddd, $J$ = 16.7, 13.7, 5.4 Hz, 1H), 2.68 – 2.47 (m, 2H), 2.02 (dp, $J$ = 10.9, 3.5 Hz, 1H), 1.60 (t, $J$ = 7.0 Hz, 2H), 1.54 (t, $J$ = 6.9 Hz, 2H), 1.36 (d, $J$ = 4.3 Hz, 6H). HRMS (ESI-MS) calcd for $C_{42}H_{39}FN_9O_5^+$ m/z $(M+H)^+$ 768.3053, found: 768.3055.

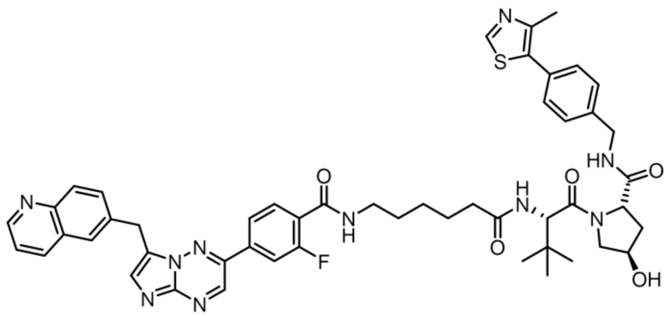

(2S,4R)-1-((S)-2-(6-(2-fluoro-4-(7-(quinolin-6-ylmethyl)imidazo[1,2-b][1,2,4]triazin-2-yl)benzamido)hexanamido)-3,3-dimethylbutanoyl)-4-hydroxy-N-(4-(4-methylthiazol-5-yl)benzyl)pyrrolidine-2-carboxamide (50-207)

Yield = 68%. $^1$H NMR (499 MHz, DMSO-$d_6$) δ 9.25 (s, 1H), 9.04 – 8.95 (m, 2H), 8.64 – 8.53 (m, 2H), 8.47 – 8.40 (m, 1H), 8.12 – 8.00 (m, 5H), 7.95 (dd, *J* = 8.7, 2.0 Hz, 1H), 7.87 (d, *J* = 9.3 Hz, 1H), 7.77 (t, *J* = 7.7 Hz, 1H), 7.72 (dd, *J* = 8.3, 4.6 Hz, 1H), 7.46 – 7.36 (m, 4H), 4.69 (s, 2H), 4.56 (d, *J* = 9.4 Hz, 1H), 4.48 – 4.40 (m, 2H), 4.38 – 4.32 (m, 1H), 4.22 (dd, *J* = 15.8, 5.5 Hz, 1H), 3.73 – 3.62 (m, 2H), 3.26 (q, *J* = 6.8 Hz, 2H), 2.44 (s, 3H), 2.29 (dt, *J* = 14.7, 7.6 Hz, 1H), 2.15 (dq, *J* = 14.3, 7.5, 6.8 Hz, 1H), 2.08 – 2.00 (m, 1H), 1.91 (ddd, *J* = 12.8, 8.5, 4.6 Hz, 1H), 1.53 (qt, *J* = 11.9, 6.8 Hz, 4H), 1.37 – 1.20 (m, 2H), 0.94 (s, 9H). HRMS (ESI-MS) calcd for $C_{50}H_{54}FN_{10}O_5S^+$ m/z $(M+H)^+$ 925.3978, found: 925.3977.

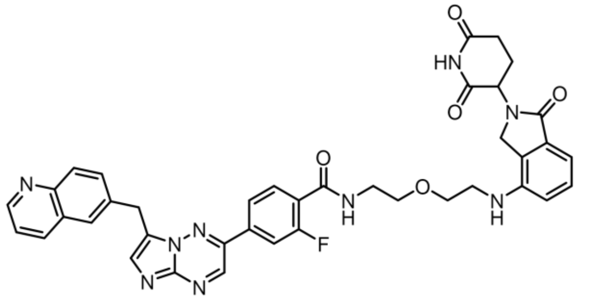

N-(2-(2-((2-(2,6-dioxopiperidin-3-yl)-1-oxoisoindolin-4-yl)amino)ethoxy)ethyl)-2-fluoro-4-(7-(quinolin-6-ylmethyl)imidazo[1,2-b][1,2,4]triazin-2-yl)benzamide (50-208)

Yield = 78%. $^1$H NMR (499 MHz, DMSO-$d_6$) δ 10.99 (s, 1H), 9.24 (s, 1H), 8.99 (dd, *J* = 4.6, 1.6 Hz, 1H), 8.58 (d, *J* = 8.4 Hz, 1H), 8.47 (dt, *J* = 5.8, 3.7 Hz, 1H), 8.11 – 8.03 (m, 2H), 8.02 (dd, *J* = 7.8, 2.1 Hz, 3H), 7.94 (dd, *J* = 8.8, 2.0 Hz, 1H), 7.77 (t, *J* = 7.7 Hz, 1H), 7.69 (dd, *J* = 8.4, 4.6 Hz, 1H), 7.28 (t, *J* = 7.7 Hz, 1H), 6.93 (d, *J* = 7.4 Hz, 1H), 6.82 (d, *J* = 8.1 Hz, 1H), 5.09 (dd, *J* = 13.3, 5.1 Hz, 1H), 4.70 (s, 2H), 4.22 (d, *J* = 17.1 Hz, 1H), 4.12 (d, *J* = 17.1 Hz, 1H), 3.66 (t, *J* = 5.8 Hz, 2H), 3.60 (t, *J* = 5.8 Hz, 2H), 3.47 (q, *J* = 5.8 Hz, 2H), 3.34 (t, *J* = 5.8 Hz, 2H), 2.89 (ddd, *J* = 17.2, 13.6, 5.5 Hz, 1H), 2.55 (s, 1H), 2.24 (qd, *J* = 13.2, 4.5 Hz, 1H), 2.00 (dtd, *J* = 11.8, 6.7, 6.0, 3.3 Hz, 1H). HRMS (ESI-MS) calcd for $C_{39}H_{35}FN_9O_5^+$ m/z $(M+H)^+$ 728.2740, found: 728.2741.

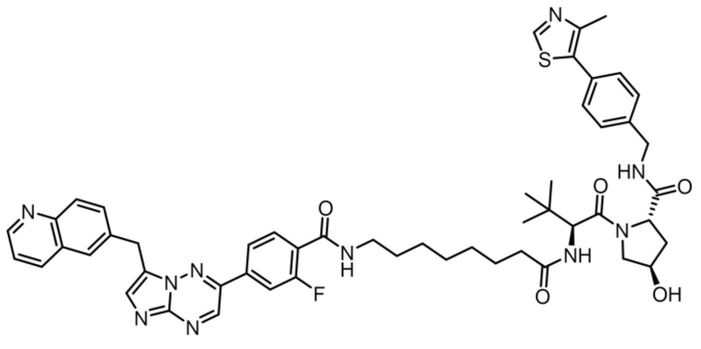

(2S,4R)-1-((S)-2-(8-(2-fluoro-4-(7-(quinolin-6-ylmethyl)imidazo[1,2-b][1,2,4]triazin-2-yl)benzamido)octanamido)-3,3-dimethylbutanoyl)-4-hydroxy-N-(4-(4-methylthiazol-5-yl)benzyl)pyrrolidine-2-carboxamide (50-209)

Yield = 71%. $^{1}$H NMR (499 MHz, DMSO-$d_6$) δ 9.25 (s, 1H), 9.03 – 8.95 (m, 2H), 8.63 – 8.53 (m, 2H), 8.47 – 8.41 (m, 1H), 8.12 – 8.01 (m, 5H), 7.94 (dd, *J* = 8.8, 2.0 Hz, 1H), 7.85 (d, *J* = 9.3 Hz, 1H), 7.76 (t, *J* = 7.7 Hz, 1H), 7.71 (dd, *J* = 8.3, 4.6 Hz, 1H), 7.46 – 7.35 (m, 4H), 4.69 (s, 2H), 4.55 (d, *J* = 9.4 Hz, 1H), 4.48 – 4.39 (m, 2H), 4.35 (s, 1H), 4.22 (dd, *J* = 15.9, 5.4 Hz, 1H), 3.71 – 3.61 (m, 4H), 3.26 (q, *J* = 6.7 Hz, 2H), 2.44 (s, 3H), 2.28 (dt, *J* = 14.8, 7.7 Hz, 1H), 2.13 (dt, *J* = 14.2, 7.1 Hz, 1H), 2.03 (t, *J* = 10.7 Hz, 1H), 1.91 (ddd, *J* = 12.9, 8.6, 4.6 Hz, 1H), 1.50 (d, *J* = 15.9 Hz, 4H), 1.31 (d, *J* = 6.9 Hz, 5H), 0.94 (s, 9H). HRMS (ESI-MS) calcd for $C_{52}H_{58}FN_{10}O_5S^+$ m/z $(M+H)^+$ 953.4291, found: 953.4293.

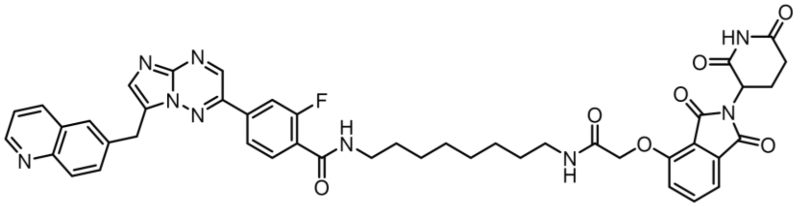


N-(8-(2-((2-(2,6-dioxopiperidin-3-yl)-1,3-dioxoisoindolin-4-yl)oxy)acetamido)octyl)-2-fluoro-4-(7-(quinolin-6-ylmethyl)imidazo[1,2-b][1,2,4]triazin-2-yl)benzamide (50-210)

Yield = 70%. $^{1}$H NMR (499 MHz, DMSO-$d_6$) δ 11.12 (s, 1H), 9.24 (s, 1H), 8.98 (dd, *J* = 4.6, 1.7 Hz, 1H), 8.56 (d, *J* = 8.5 Hz, 1H), 8.43 (dt, *J* = 5.7, 3.3 Hz, 1H), 8.08 – 8.01 (m, 5H), 7.97 – 7.89 (m, 2H), 7.84 – 7.73 (m, 2H), 7.68 (dd, *J* = 8.4, 4.5 Hz, 1H), 7.49 (d, *J* = 7.3 Hz, 1H), 7.39 (d, *J* = 8.6 Hz, 1H), 5.12 (dd, *J* = 12.8, 5.5 Hz, 1H), 4.77 (s, 2H), 4.69 (s, 2H), 3.26 (q, *J* = 6.7 Hz, 2H), 3.15 (q, *J* = 6.7 Hz, 2H), 2.90 (ddd, *J* = 16.0, 13.6, 5.4 Hz, 1H), 2.58 (d, *J* = 35.4 Hz, 2H), 2.04 (dp, *J* = 11.6, 4.2, 3.8 Hz, 1H), 1.49 (dt, *J* = 36.4, 7.0 Hz, 5H), 1.29 (s, 7H). HRMS (ESI-MS) calcd for $C_{45}H_{43}FN_9O_7^+$ m/z $(M+H)^+$ 840.3264, found: 840.3263.

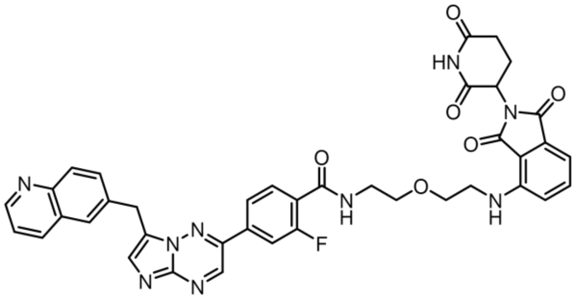


N-(2-(2-((2-(2,6-dioxopiperidin-3-yl)-1,3-dioxoisoindolin-4-yl)amino)ethoxy)ethyl)-2-fluoro-4-(7-(quinolin-6-ylmethyl)imidazo[1,2-b][1,2,4]triazin-2-yl)benzamide (50-212)

Yield = 64%. $^{1}$H NMR (499 MHz, DMSO-$d_6$) δ 11.09 (s, 1H), 9.24 (s, 1H), 8.98 (dd, *J* = 4.6, 1.7 Hz, 1H), 8.56 (d, *J* = 8.4 Hz, 1H), 8.43 (td, *J* = 5.7, 2.3 Hz, 1H), 8.10 – 7.99 (m, 5H), 7.92 (dd, *J* = 8.7, 2.0 Hz, 1H), 7.78 (t, *J* = 7.8 Hz, 1H), 7.68 (dd, *J* = 8.4, 4.6 Hz, 1H), 7.57 (dd, *J* = 8.5, 7.1 Hz, 1H), 7.17 (d, *J* = 8.6 Hz, 1H), 7.02 (d, *J* = 7.0 Hz, 1H), 6.64 (s, 1H), 5.03 (dd, *J* = 12.8, 5.4 Hz, 1H), 4.69 (s, 2H), 3.64 (dt, *J* = 30.4, 5.6 Hz, 4H), 3.49 (dq, *J* = 17.5, 5.6 Hz, 4H), 2.92 – 2.80 (m, 1H), 2.56 – 2.52 (m, 2H), 2.00 (ddd, *J* = 11.0, 6.0, 3.4 Hz, 1H). HRMS (ESI-MS) calcd for $C_{39}H_{33}FN_9O_6^+$ m/z $(M+H)^+$ 742.2532, found: 742.2532.

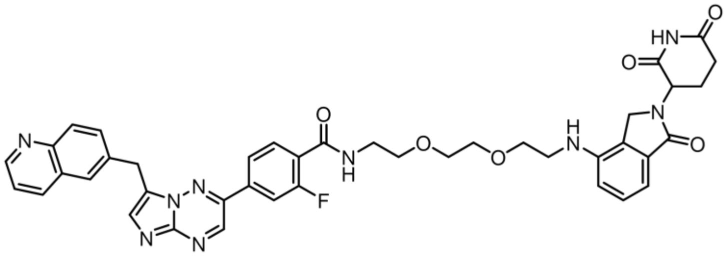

N-(2-(2-(2-((2-(2,6-dioxopiperidin-3-yl)-1-oxoisoindolin-4-yl)amino)ethoxy)ethoxy)ethyl)-2-fluoro-4-(7-(quinolin-6-ylmethyl)imidazo[1,2-b][1,2,4]triazin-2-yl)benzamide (50-213)
Yield = 42%. $^{1}$H NMR (499 MHz, DMSO-$d_6$) δ 11.00 (s, 1H), 9.23 (s, 1H), 9.00 (dd, $J$ = 4.7, 1.6 Hz, 1H), 8.59 (d, $J$ = 8.4 Hz, 1H), 8.45 (td, $J$ = 5.6, 2.3 Hz, 1H), 8.13 – 7.99 (m, 6H), 7.94 (dd, $J$ = 8.7, 2.0 Hz, 1H), 7.79 (t, $J$ = 7.9 Hz, 1H), 7.70 (dd, $J$ = 8.4, 4.6 Hz, 1H), 7.24 (t, $J$ = 7.7 Hz, 1H), 6.90 (d, $J$ = 7.4 Hz, 1H), 6.77 (d, $J$ = 8.1 Hz, 1H), 5.11 (dd, $J$ = 13.3, 5.1 Hz, 1H), 4.69 (s, 2H), 4.21 (d, $J$ = 17.1 Hz, 1H), 4.12 (d, $J$ = 17.1 Hz, 1H), 3.65 – 3.54 (m, 7H), 3.45 (p, $J$ = 6.1, 5.5 Hz, 2H), 3.31 (t, $J$ = 5.9 Hz, 2H), 2.98 – 2.86 (m, 1H), 2.69 – 2.52 (m, 2H), 2.30 (qd, $J$ = 13.2, 4.5 Hz, 1H), 2.03 (dtd, $J$ = 11.4, 6.4, 5.8, 3.1 Hz, 1H). HRMS (ESI-MS) calcd for $C_{41}H_{39}FN_9O_6$+ m/z $(M+H)^+$ 772.3002, found: 772.3000.

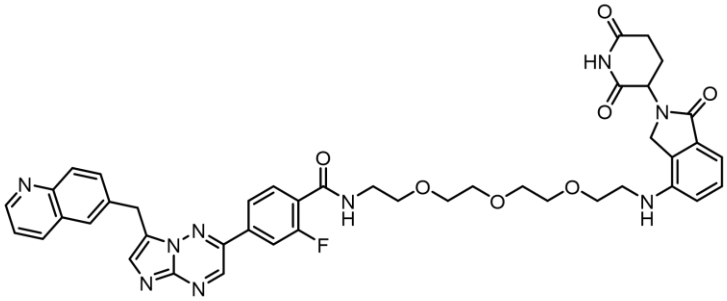

N-(2-(2-(2-(2-((2-(2,6-dioxopiperidin-3-yl)-1-oxoisoindolin-4-yl)amino)ethoxy)ethoxy)ethoxy)ethyl)-2-fluoro-4-(7-(quinolin-6-ylmethyl)imidazo[1,2-b][1,2,4]triazin-2-yl)benzamide (50-214)
Yield = 62%. $^{1}$H NMR (499 MHz, DMSO-$d_6$) δ 11.00 (s, 1H), 9.24 (s, 1H), 8.99 (dd, $J$ = 4.6, 1.6 Hz, 1H), 8.57 (d, $J$ = 8.4 Hz, 1H), 8.45 (td, $J$ = 5.6, 2.3 Hz, 1H), 8.10 – 8.00 (m, 6H), 7.93 (dd, $J$ = 8.8, 2.0 Hz, 1H), 7.80 (t, $J$ = 7.8 Hz, 1H), 7.69 (dd, $J$ = 8.4, 4.6 Hz, 1H), 7.24 (t, $J$ = 7.7 Hz, 1H), 6.91 (d, $J$ = 7.4 Hz, 1H), 6.77 (d, $J$ = 8.1 Hz, 1H), 5.11 (dd, $J$ = 13.3, 5.1 Hz, 1H), 4.68 (s, 2H), 4.21 (d, $J$ = 17.1 Hz, 1H), 4.11 (d, $J$ = 17.1 Hz, 1H), 3.62 – 3.52 (m, 4H), 3.55 (s, 8H), 3.44 (q, $J$ = 5.9 Hz, 2H), 3.30 (t, $J$ = 6.0 Hz, 2H), 2.92 (ddd, $J$ = 17.5, 13.7, 5.5 Hz, 1H), 2.65 – 2.58 (m, 1H), 2.30 (qd, $J$ = 13.2, 4.5 Hz, 1H), 2.03 (dtd, $J$ = 11.4, 6.3, 5.8, 3.1 Hz, 1H). HRMS (ESI-MS) calcd for $C_{43}H_{43}FN_9O_7^+$ m/z $(M+H)^+$ 816.3264, found: 816.3265.

**Molecular modeling and docking analysis**

Scrödinger small molecule drug discovery suite 2020–1 was used for Molecular modeling. The crystal structure of c-MET kinase (pdb:3zbx) was retrieved from the protein data bank. The protein structure was analyzed using Maestro version 12.3.013 (Scrödinger Inc.) and subjected to docking. The Protein Preparation Wizard was used to add missing hydrogen atoms and side chains and minimized using OPLS3e force field to optimize hydrogen bonding network and converge the heavy atoms to a rmsd of 0.3 Å. The

Capmatinib structure was drawn in Maestro and subjected to Lig Prep to generate conformers, possible protonation at pH of 7 ± 2 that serves as an input for docking process. The receptor grid was generated using Receptor Grid Generation tool in Maestro (Scrödinger Inc.). Docking was performed using GLIDE XP with the van der Waals radii of nonpolar atoms for each of the ligands were scaled by a factor of 0.8 and partial charge cutoff 0.15. The binding mode of the ligand in the protein was analyzed using Maestro version 12.3.013 (Scrödinger Inc.).

**Fluorescent models**

HEK293T (from Mayo Clinic's existing stock) cells were cultured in Dulbecco's Modified Eagle's Medium (DMEM, Corning), 10% Fetal Bovine Serum (FBS, Corning)), Penicillin/Streptomycin (1:100, Gibco). The plasmid pLenti-MetGFP (Addgene #37560) and the exon 14 skipping mutation was introduced using Q5 Site-Directed Mutagenesis Kit (NEB, Cat# E0554S). In brief, we used 25ng of pLenti-MET-GFP plasmid as a template, 10μM forward primer(5'-ATCAGTTTCCTAATTCATTCTAG-3') and 10μM reverse primer(5'-CTTTAATTTGCTTTCTCTTTTTC-3'), 12.5ul of Q5 Hot Start High-Fidelity 2X master Mix for mutagenesis PCR (30 seconds in 98$^{o}$C for initial denaturation, 98$^{o}$C for 10 seconds, 62$^{o}$C for 30 seconds and 72$^{o}$C for 6 minutes, repeated for 25 cycles, plus 2 minutes in 72$^{o}$C for final extension). The PCR product was processed with KLD reaction and transformation following the protocol provided by the manufacturer (NEB Inc.).

***MET*ex14Δ-GFP screening**

HEK293 *MET*ex14Δ expressing cells (4 x $10^3$ cells / well) were plated in flat bottomed, sterile, 96 well TPP tissue culture plates (Catalog no. 92696 Midwest Scientific), in 90 µL DMEM high media (Catalog no. SH30022 VWR). The cells were allowed to adhere overnight at 37°C in a $CO_2$ incubator (5% $CO_2$). Drug working solutions were prepared at 10 mM in DMSO (Catalog no. BP231 Fisher Scientific) stocks. 10 µL of the working solution was added to each well in the plate to yield the indicated final concentration in 0.1% DMSO. A DMSO control was included. The plates were set up in an Incucyte SX3 live cell imager (Sartorius Corporation) to acquire phase contrast and green fluorescence data. The imager was itself housed in a $CO_2$ incubator (5% $CO_2$) and the plates were maintained at 37°C over the course of the experiment. Images of the entire well were captured at 4x magnification, every 2 hours, for the indicated period. The images were analyzed with the included phase contrast cell tracking module software to generate the curves showing % confluence over time. The degradation percentage was calculated using the following formula: Percent Degradation = 100 – (100x ($F_x/F_{DMSO}$)) where $F_x$ is the normalized green count in the treatment groups and $F_{DMSO}$ is the normalized green count in the DMSO control group.

**Sample preparation for proteomics**

Suspensions of 5 million cells (HEK293 *MET*ex14Δ-GFP) were seeded into 15 cm plates in 20 mL of DMEM high (catalog no. SH30021.FS Hyclone) with 10% FBS (Catalog no. 26140079 Life Technologies) and 1% Penicillin- Streptomycin (16777-164, Hyclone) and were allowed to adhere overnight in a humidified 5% $CO_2$ incubator at 37°C. The following day, cells were treated with 1 µM of capmatinib, 1 µM of 48-284, or DMSO and incubated in a humidified 5% $CO_2$ incubator at 37°C for 24 hours. Cell plates were washed twice with cold 1 × PBS, harvested in cold 1 × PBS, and centrifuged at 14,000 g for 10 min at 4°C. Cells were lysed in Radioimmunoprecipitation assay (RIPA) buffer (Thermo Scientific), sodium orthovandate ($Na_3VO_4$, Sigma), sodium fluoride (NaF, Sigma), β-glycerophosphate (Sigma) and 1 mM phenylmethylsulfonyl fluoride (PMSF, Sigma). The samples were incubated on ice for 30 minutes, vortexed at 10 min intervals and centrifuged at 14,000 g for 10 minutes at 4°C. The supernatant was collected, and protein quantified using the BCA Protein Assay kit (23225, Pierce). Samples were stored at -80°C until required for quantitative proteomics analysis. Proteins were then precipitated using cold acetone to remove detergents and the pellet washed 3 times with 70% ethanol. The proteins were dissolved in 7M urea, 2M thiourea, 5 mM DTT, 100mM tris/HCl, pH 7.8. Protein amounts were quantified using the CB-X™ protein assay (G-Biosciences, St Louis, MO) and a 50 ug aliquot of each sample was alkylated with iodoacetamide (final concentration of 20 mM) for 30 min at RT in darkness, then quenched with an equimolar amount of DTT. Samples were diluted to 1 M urea and digested with 1 µg of trypsin to give an enzyme:substrate ratio of 1:50 at 24°C overnight. Digests were acidified to 1% TFA, ~pH 3 for LC-MS/MS analysis.

**Label-free LC-MS/MS analysis**

Each sample was analyzed by LC-MS/MS on an RSLCnano system (ThermoFisher Scientific) coupled to an Orbitrap Eclipse Tribrid mass spectrometer (ThermoFisher Scientific). The samples were first injected onto a trap column (Acclaim PepMap™ 100, 75µm x 2 cm, ThermoFisher Scientific) and desalted for 3.0 minutes at a flow rate of 5 µL/min, before switching in line with the main column. Separation was performed on a C18 nano column (Acquity UPLC® M-class, Peptide CSH™ 130A, 1.7µm 75µm x 250mm, Waters Corp) at 300 nL/min with a linear gradient from 4-22% over 96 min. The LC aqueous mobile phase contained 0.1% (v/v) formic acid in water and the organic mobile phase contained 0.1% (v/v) formic acid in 100% (v/v) acetonitrile. Mass spectra for the eluted peptides were acquired in the Orbitrap using the data-dependent mode with a mass range of m/z 250–1500, resolution 120,000, AGC target 4 x $10^6$, maximum injection time 50 ms for the MS1 peptide measurements. Data-dependent MS2 spectra were acquired by HCD in the ion trap with a normalized collision energy (NCE) set at 30%, AGC target set to 1 x $10^5$, intensity threshold 1 x $10^5$ and a maximum injection time of 35 ms. Dynamic exclusion was set at 45 sec and the isolation window set to 1.6 m/z.

**Proteomics data analysis**

The identification and quantitation of the proteins were performed using Proteome Discoverer (Thermo; version 2.4). All MS/MS samples were searched using Mascot (Matrix Science, London, UK; version 2.6.2). Mascot was set up to search an in-house modified version of the cRAP_20150130.fasta database (124 entries); uniprot-human_20201207 database (75777 entries) assuming the digestion enzyme trypsin and a maximum of 2 missed cleavages. Mascot was searched with a fragment ion mass tolerance of 0.6 Da and a parent ion tolerance of 10.0 ppm. Deamidated of asparagine and glutamine, oxidation of methionine, and protein N-terminal acetylation were specified in Mascot as variable modifications, while carbamidomethyl of cysteine was fixed. Peptides were validated by Percolator with a 0.01 posterior error probability (PEP) threshold. The data were searched using a decoy database to set the false discovery rate to 1% (high confidence). Only proteins identified with a minimum of 2 unique peptides were further analyzed for quantitative changes. The peptides were quantified using the precursor abundance based on intensity. The peak abundance was normalized using total peptide amount. The peptide group abundances are summed for each sample and the maximum sum for all files is determined. The normalization factor used is the factor of the sum of the sample and the maximum sum in all files. The protein ratios are calculated using the summed abundance for each replicate separately, and the geometric median of the resulting ratios is used as the protein ratios. The significance of differential expression was tested using an ANOVA test which provides a p-value and an adjusted p-value using the Benjamini-Hochberg method for all the calculated ratios. The mass spectrometry proteomics data have been deposited to the ProteomeXchange Consortium via the PRIDE [31] partner repository with the dataset identifier PXD031614.

**Western blots**

The Hs746T and H596 cell lines were purchased from the American Type Culture Collection. These cells were lysed after the specified treatments in an appropriate volume of NETN lysis buffer (5M NaCl, 0.5M EDTA, PH8.0, 1M Tris-HCl, PH8.0, NP-40-0.5%) with a protease inhibitor cocktail tablet (complete Mini, EDTA-free, Roche Cat#11836170001). Protein concentration was measured using Pierce BCA protein assay kit (Thermo Fisher Scientific) and adjusted to 10 to 50 mg per line, using 5X Laemmli Sample Buffer with the addition of 10% 2-mercaptoethanol (Sigma Aldrich). Samples were run on 4 - 20% Mini-Protean TGX pre-cast protein gels and transferred to PVDF membranes using the Trans-Blot Turbo Transfer System (all Bio-Rad). Membranes were blocked for 1 hour at room temperature using 4% milk in TBS-T (20 mM Tris-HCl pH 7.5, 150 mM NaCl, 0.1% Tween20), then membranes were incubated in primary antibody at 4°C overnight. After washing three times, membranes were incubated with secondary antibody at room temperature for 1 hour. For imaging, the membranes were incubated in SuperSignal West Pico Plus Chemiluminescent Substrate (Thermo scientific) and developed with HyBlot Cl autoradiography film and X-omat 2000A system (Kodak). The films were scanned by Epson Perfection 2400Photo. ImageJ was used to quantify bands in the Western blot, normalized to the Beta-actin control for each lane.

**Antibodies**

The following antibodies were used for the Western blots described above:
Anti-MET Cell Signaling Technology (Cat#4060)
Anti-Phosphorylated-MET(Y1234/1235): Cell Signaling Technology (Cat#3077S)
Anti-Akt: Cell Signaling Technology (Cat#4691P)
Anti-Phosphorylation Akt (Ser473) Cell Signaling Technology (Cat#4060)
Anti-p44/42 MAPK(Erk1/2): Cell Signaling Technology (Cat#4695)
Anti-Phosphorylation p44/42 MAPK(Erk1/2)9T202/Y204): Cell Signaling Technology (Cat#4370S)
Anti-Ubiquitin(P37): Cell Signaling Technology (Cat#58395S)
Anti-Beta-Actin: Santa Cruz Biotechnology (SC-47778)
Anti-GAPDH: Santa Cruz Biotechnology (SC-32233))
Anti-GFP Cell Signaling Technology (Cat#2555S)

**Short perturbation experiment and MET quantification**

We performed a short-term treatment experiment on the UW21 model to determine the immediate effects of 48-284 on MET. The tumor xenograft model UW21 was provided to our team by Andrew Baschnagel, MD at the University of Wisconsin through a materials transfer agreement. The UW21 xenograft was derived from the brain metastasis of a patient with non-small cell lung cancer and contains a *MET*ex14Δ mutation as characterized previously [23]. 48-284 was dissolved in solvent (5% DMSO, 5% Solutol, Kollipgor HS15, Sigma-Aldrich, Cat#42966) at 5mg/ml concentration and 15mg/kg body weight was administered twice through mouse tail vein injection at time point 0 hours and 8 hours. Six hours after the second dose, PDX tumor tissues were harvested, one piece was snap frozen and one was fixed in 10% formalin. Xenograft experiments in mice were performed under a protocol approved by the Mayo Clinic's Institutional Animal Care and Use Committee. Following treatment with 48-284, the UW21 xenograft was removed from the flanks of the mice, fixed in formalin, embedded in paraffin, sectioned at 4 microns and stained with an antibody directed against MET (clone SP44, Ventana, Tucson, AZ, USA). ImageJ was used to quantify positively stained cells from both the treated and untreated xenografts. We compared MET staining in multiple regions on interest in both the treated and untreated xenografts. We also performed a Western blot as per above. ImageJ was used to quantify bands in the Western blots, normalized to the beta-actin control for each lane. We compared the results between treatment groups with a two-way, unpaired t test using R and the tidyverse packages and considered p values $< 0.05$ significant.

Please note: Animal experiments were performed as described in protocol IACUC A00006039 that was approved by IACUC committee Mayo Clinic, Rochester MN. All the experiments were performed in compliance with relevant laws or guidelines.

**Raw data**

### Figure 1B (raw images)

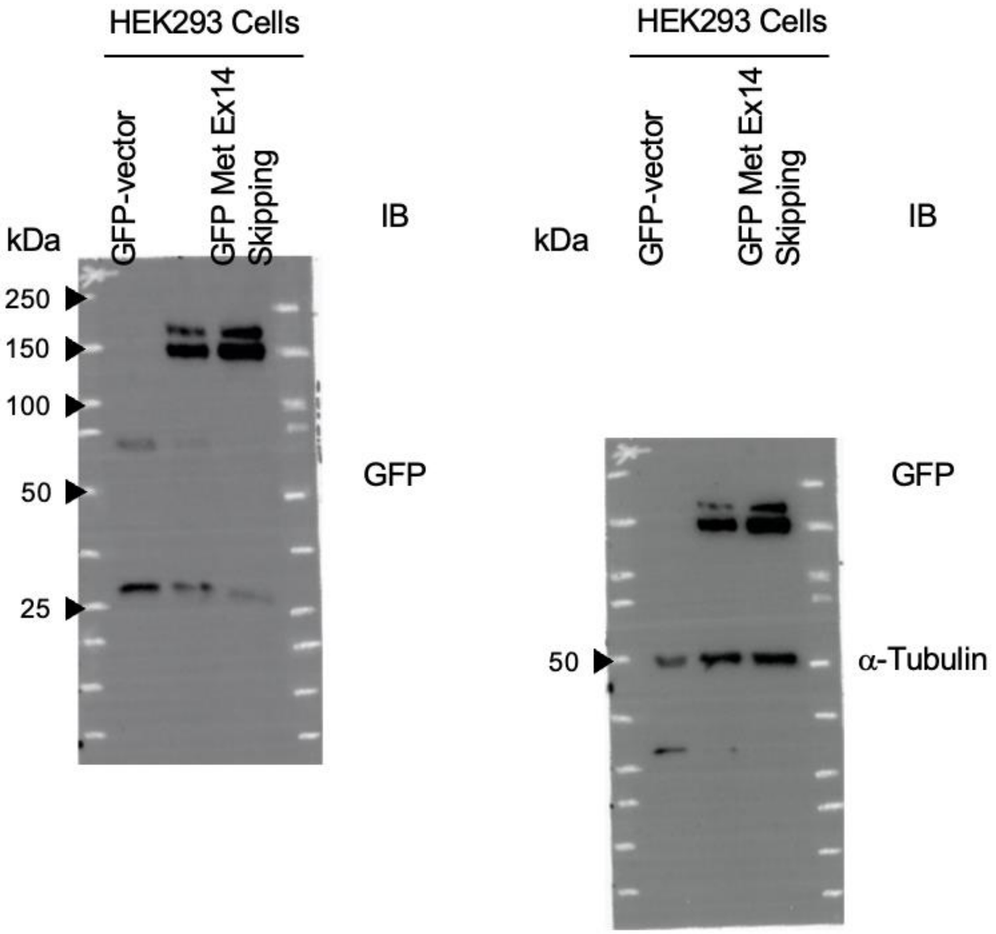

Figure 3 A (raw images)

Hs746T-48-284

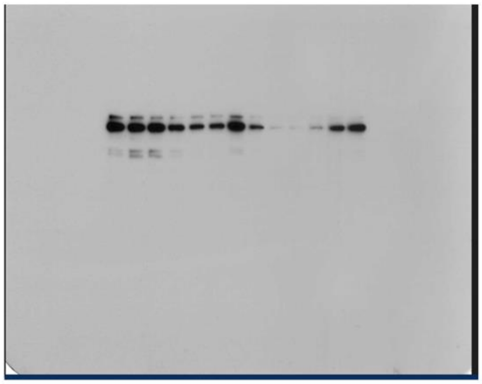


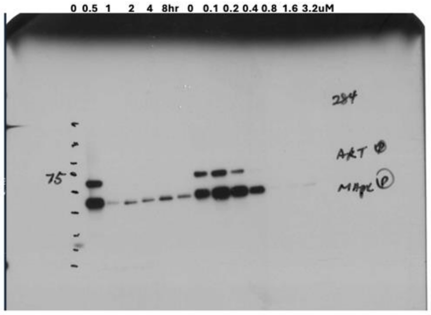


c-MET

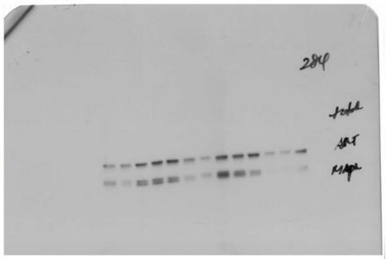


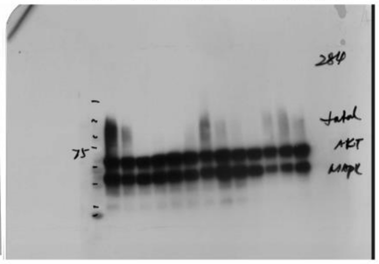


Figure 3 B (raw images)

Hs746T-SJF8240

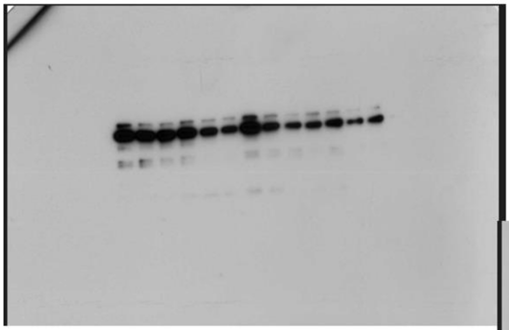


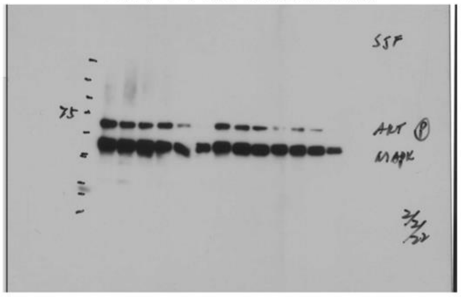


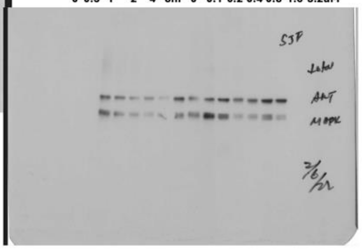


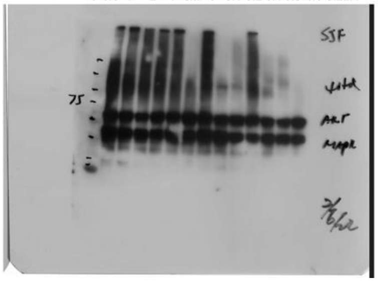

Figure 3 C (raw images)

PDX model-48-284

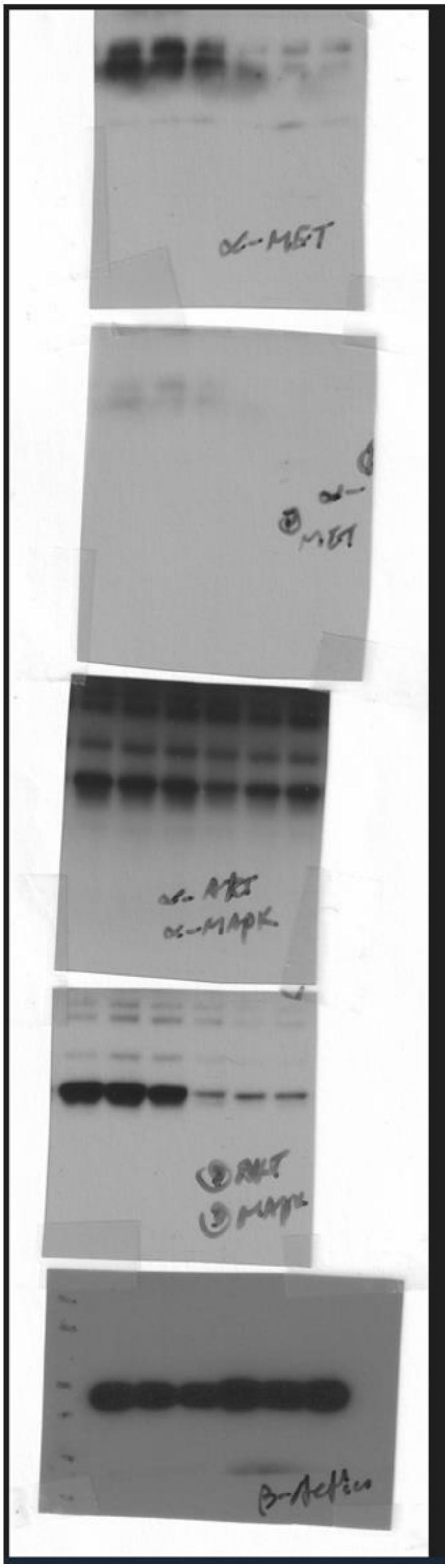

Figure S3 (raw images)

H596-48-284

Timepoints(hours) Doses(uM)

0 0.5 1 2 4 8hr 0 0.1 0.2 0.4 0.8 1.6 3.2uM

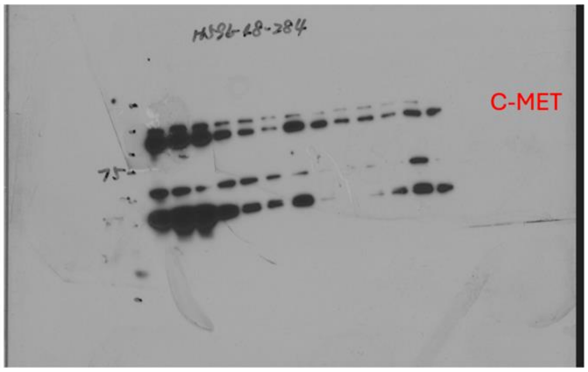


Timepoints(hours) Doses(uM)

0 0.5 1 2 4 8hr 0 0.1 0.2 0.4 0.8 1.6 3.2uM

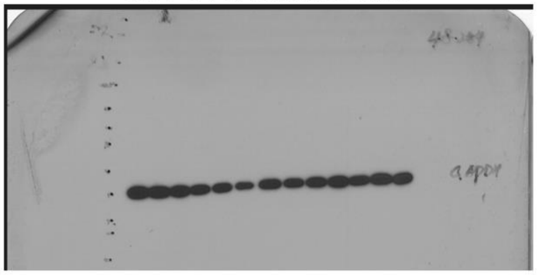
